# IgA Targeting in the Infant Gut Is Modulated by Diet and Increasingly Directed Towards Persistent Species

**DOI:** 10.64898/2026.05.19.726352

**Authors:** Jing Qian, Parsa Ghadermazi, Soren Maret, Jennifer F. Kemp, Daniel Frank, Edward L. Melanson, Audrey E. Hendricks, Nancy Krebs, Minghua Tang, Matthew R. Olm

## Abstract

**Background:** IgA is the dominant antibody in the human gut and a key regulator of host-microbe interactions. Infants begin to produce IgA at around 6 months old and receive large quantities of IgA via human milk, but technical limitations have prevented species-level characterization of IgA binding in early life. This has left basic knowledge gaps about which species are targeted by IgA in infancy, and how modifiable lifestyle factors like breastfeeding and complementary feeding impact IgA targeting.

**Results:** Here we adapt Metagenomic Immunoglobulin Sequencing (MIg-Seq) for low-biomass infant fecal samples and apply this optimized protocol to 32 longitudinal samples from 16 infants enrolled in the MINT trial, a four-arm randomized controlled trial comparing meat-based, dairy-based, plant-based, and reference complementary feeding patterns, with fecal sampling at 6 and 12 months (pre and post intervention). Infant IgA targeting mirrors adults at the phylum level, with both age groups showing significantly higher IgA targeting of Pseudomonadota and lower targeting of Bacteroidota relative to other phyla. During the substantial microbiome compositional shifts noted between 6 and 12 months, IgA targeting is significantly more stable than the microbiome itself. Among persistent colonizers, IgA targeting strengthens significantly from 6 to 12 months, with the most pronounced effect observed for *Bifidobacterium*, a finding robust across all dietary arms and feeding modes. The feeding arm to which infants were enrolled was not significantly associated with IgA binding, but several nutrient-specific associations were discovered. Animal-derived nutrients, particularly cholesterol, are strongly positively correlated with IgA targeting of *Bifidobacterium longum*, while plant-derived carotenoids are positively associated with IgA targeting of *Flavonifractor plautii* and *Ruminococcus gnavus*.

**Conclusions:** This study introduces an experimental and computational framework for species-level IgA profiling in the infant gut. The progressive strengthening of IgA targeting of *Bifidobacterium* and other beneficial persistent colonizers suggests a role for IgA in reinforcing beneficial microbes during infancy. The nutrient-specific dietary effects on IgA targeting reveal the immunological consequences of the complementary feeding period, and highlight a contrast between animal-versus plant-based diets. Together, these findings point to early nutritional interventions and IgA-based therapeutics as promising tools for promoting healthy immune-microbiome development.

## Background

The human gut undergoes a dramatic transformation during the first year of life, expanding from a sterile or near-sterile environment at birth to a complex, stable community of hundreds to thousands of bacterial species by the end of infancy. During this window, the developing mucosal immune system must establish tolerance to the colonizing commensal community and maintain responsiveness to pathogens. Disruptions to early microbial colonization have been associated with increased risks of allergic disease, asthma, inflammatory bowel disease, and metabolic disorders ^1^, conditions whose incidence has risen sharply in industrialized societies over recent decades ^2,3^. How the developing immune system monitors and responds to this rapidly changing microbiota, and how early-life exposures such as diet shape this process, remains unclear.

Immunoglobulin A (IgA) is the most abundantly secreted antibody in the human gut and serves as a primary mediator of immune-microbiota homeostasis. Through selective coating of bacterial taxa, IgA influences which members of the colonizing microbiota are recognized and retained, making the pattern of IgA targeting across the community a functional readout of the host-microbe relationship ^4,5^. IgA binding can have opposing consequences for its targets: IgA can restrict bacterial growth by enchaining dividing cells and blocking epithelial contact ^6^, while IgA binding can also promote stable mucosal colonization of commensals such as *Bacteroides fragilis* ^4^. However, whether IgA predominantly restricts or supports microbial taxa in the healthy infant gut remains unclear. Resolving this distinction is important for understanding how IgA shapes early microbiome assembly and for developing IgA-based strategies to modulate the infant microbiome.

During early life, gut IgA is almost entirely maternally derived. Human milk contains secretory IgA (SIgA) at concentrations of approximately 7.5 g/L in colostrum, declining to 1.6∼2 g/L in mature milk ^7^. Milk SIgA is produced by plasma cells that originate in maternal gut-associated lymphoid tissue and home to the mammary gland, and therefore reflects the mother’s microbial exposure history ^8^. Infants do not begin producing endogenous IgA-positive B cells until approximately four weeks of age ^9^, and the infant IgA repertoire remains qualitatively distinct from that of adults through at least the first year of life, with lower somatic mutation frequencies and limited evidence of antigen-driven selection ^10^. The introduction of complementary foods between approximately 6 and 12 months of age represents a major environmental transition that coincides with rapid microbiome maturation and the shift from maternally derived to endogenously produced IgA ^11–14^. Whether the type of solid food introduced during this window influences IgA targeting of the gut microbiome has not been tested.

The development of IgA sequencing approaches, including Next-generation IgA-SEQ (ng-IgA-SEQ) and Metagenomic Immunoglobulin Sequencing (MIg-seq), has made it possible to directly characterize IgA targeting at the species level by physically separating IgA-coated bacteria from the total fecal community prior to sequencing ^15,16,17^. Applied to adult cohorts, these methods have revealed that IgA targeting is highly selective, dynamically regulated, and associated with host immune status and disease ^18^. However, these methods have never been applied to infants, in part due to the low microbial biomass of infant fecal samples, leaving the species-level landscape of IgA targeting during early life uncharacterized.

Here, we adapt MIg-Seq for low-biomass infant fecal samples and apply it to a cohort of infants enrolled in a randomized dietary intervention trial comparing plant-based and meat-based, and dairy-based complementary feeding patterns. We characterize the species-level IgA-microbiome interface during infancy, compare it to established adult patterns, and provide evidence that early complementary feeding reshapes immune targeting of the gut microbiota.

## Methods

### 1. Study design and participant cohort

Full details on participant recruitment, inclusion and exclusion criteria, and the dietary intervention protocol were previously reported^19^. Briefly, infant fecal samples were collected from participants in a randomized controlled trial (the MINT study) designed to assess the impact of different complementary feeding strategies on infant growth and gut health (ClinicalTrials.gov: NCT05012930). Infants were randomized to one of four dietary intervention groups beginning at approximately 5 months of age: plant-based, meat-based, dairy-based, or a reference group receiving standard-of-care feeding guidance.

For the present study, we randomly selected 16 infants who had fecal samples available at both pre-intervention and post-intervention timepoints, yielding 32 fecal samples. Pre-intervention samples were collected between 5 and 6 months of age (prior to the start of complementary feeding), and post-intervention samples were collected at 12 months of age; all pre-intervention samples are referred to as the ‘6-month’ timepoint throughout. The 16 infants included 8 from the plant-based group, 6 from the meat-based group, and 1 each from the dairy-based and reference groups; diet-specific comparisons were restricted to the plant-based and meat-based groups. Among these infants, 7 were exclusively human milk-fed and 9 were exclusively formula-fed at the time of sampling (Table 1).

**Table 1.** Characteristics of participants included in this sub-study.

| Characteristic | Meat(n = 6) | Dairy(n = 1) | Plant(n = 8)<sup>a</sup> | Reference(n = 1) |
| --- | --- | --- | --- | --- |
| Sex: female<sup>b</sup> | 4 (67%) | 1 (100%) | 5 (71%)<sup>b</sup> | 0 (0%) |
| C-section delivery<sup>b</sup> | 2 (33%) | 0 (0%) | 3 (43%)<sup>b</sup> | 1 (100%) |
| Attending childcare<sup>b</sup> | 2 (33%) | 1 (100%) | 1 (14%)<sup>b</sup> | 0 (0%) |
| Human milk feeding | 2 (33%) | 0 (0%) | 5 (62%) | 0 (0%) |
| 12-month dietary records: |  |  |  |  |
| Complete (3 days) | 6 (100%) | 1 (100%) | 4 (50%) | 1 (100%) |
| Sparse (2 days) | 0 (0%) | 0 (0%) | 1 (12%) | 0 (0%) |
| Sparse (1 day) | 0 (0%) | 0 (0%) | 1 (12%) | 0 (0%) |
| None (0 days) | 0 (0%) | 0 (0%) | 2 (25%) | 0 (0%) |

<sup>a</sup>Clinical metadata (sex, delivery mode, and childcare attendance) were unavailable for one Plant arm infant.
<sup>b</sup>Percentage calculated among n = 7 with available data. Complementary category makes up the remainder.

All 16 infants were included in analyses characterizing general IgA targeting patterns and community structure (Figs. 2, 3, 4). Clinical metadata (sex, delivery mode, and childcare attendance) were available for 15 of 16 infants; one subject (MINT-009, Plant arm) had no recorded demographic information and was excluded from all covariate analyses, leaving n = 15 for those comparisons. Three-day dietary records collected at 12 months were available for 14 of 16 infants; two subjects (both in the Plant arm) had no dietary data and were excluded from all diet–IgA correlation analyses, leaving n = 14 for those analyses. The first diet recording day occurred a median of approximately one week after stool sample collection; given that complementary feeding practices and diet arm assignment were sustained throughout the intervention period, these records are considered representative of dietary intake patterns contemporaneous with microbiome sampling. Among the 14 infants with dietary records, 12 had a complete full three-day assessment. The other 2 had days of complete recordings, and were retained in dietary analyses. Daily nutrient intakes were calculated as the sum of recorded intakes divided by the number of available days. Demographic and clinical characteristics are summarized in Table 1 (See Results).

**Figure 1.**
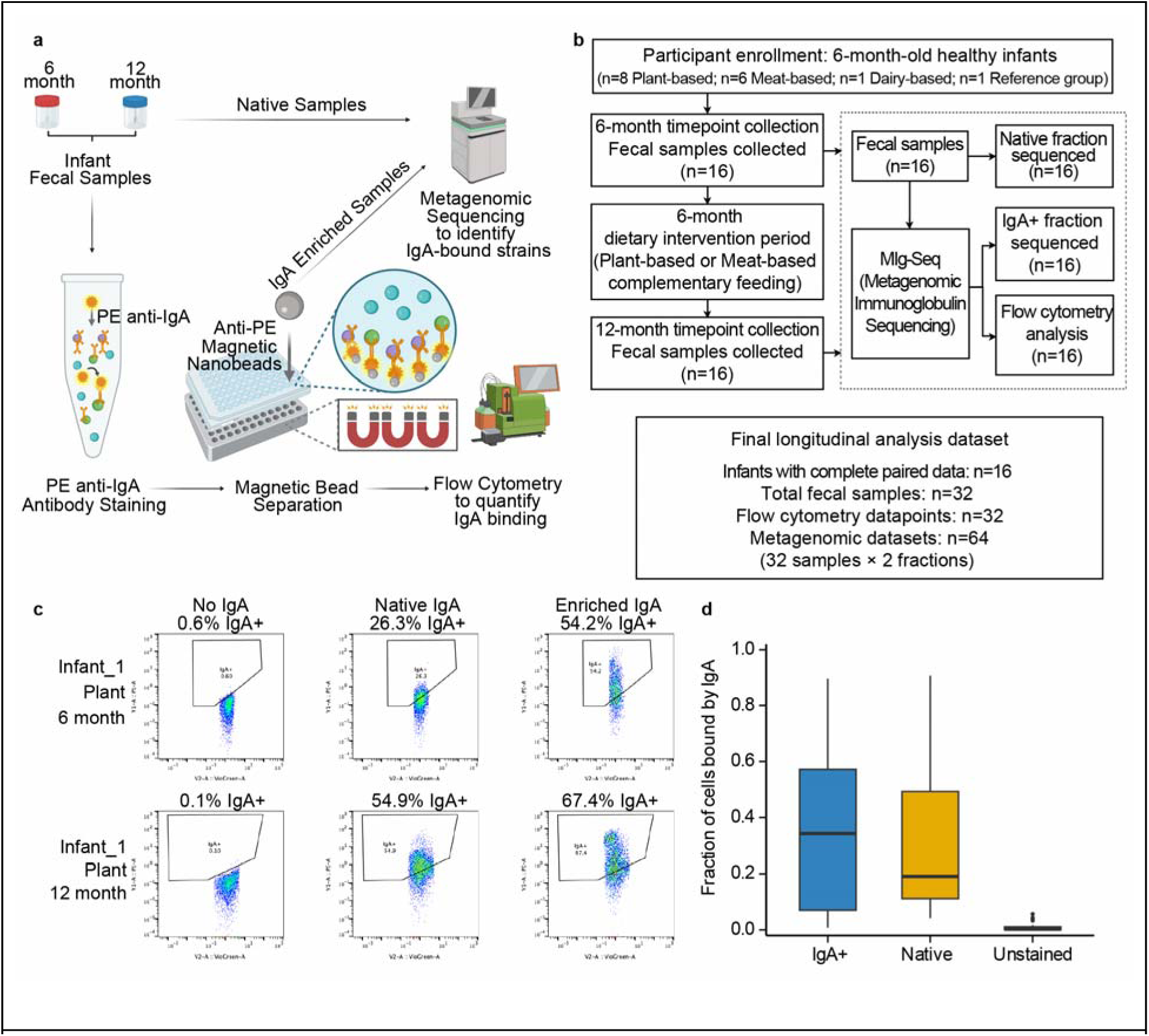
Adaptation and validation of the MIg-Seq workflow for infant fecal samples. **a**, Schematic overview of the MIg-Seq workflow adapted for low-biomass infant fecal samples. **b**, Study design and data generation flowchart. **c**, Representative flow cytometry plots from a representative infant at 6 months (top) and 12 months (bottom), showing unstained control (left), native sample (middle) and magnetically enriched IgA+ fraction (right). Bacterial cells were identified by SYBR Green fluorescence (x-axis) and IgA-coated cells by PE anti-IgA signal (y-axis). **d**, Fraction of total microbial cells bound by IgA, as measured by flow cytometry, in enriched IgA+, native, and unstained fractions. n = 32. Boxplots show the median (centre line), interquartile range (box bounds).

**Figure 2.**
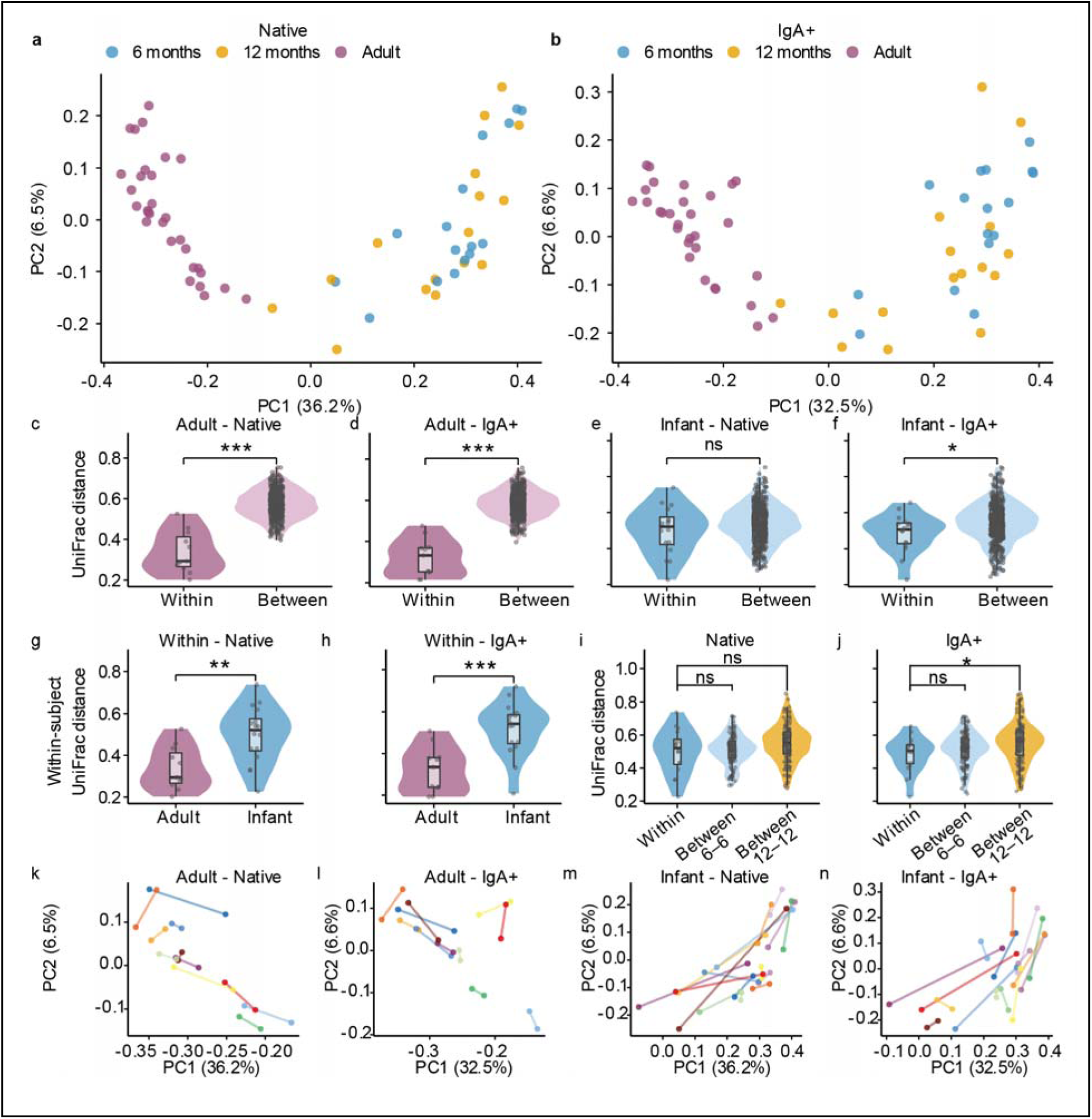
β-diversity analysis reveals age-driven community structure and progressive individual specificity in the IgA-coated infant microbiome. **a**, **b**, PCoA of unweighted UniFrac distances for native **(a)** and IgA+ **(b)** fractions. Points colored by age group (6-month infant, 12-month infant, Adult). Pairwise PERMANOVA (Infant vs Adult): native R² = 0.329, IgA+ R² = 0.285; *p* = 0.001 for both. n = 62 samples (32 infant, 30 adult). **c–f**, Within-subject versus between-subject UniFrac distances for adults (**c**, native; **d**, IgA+) and infants (**e**, native; **f**, IgA+). Wilcoxon rank-sum test. **g, h**, Within-subject temporal distances compared between adults and infants for native **(g)** and IgA+ **(h)** fractions. Wilcoxon rank-sum test. **i, j**, Between-subject distances in infants stratified by timepoint pair (6mo–6mo, 12mo–12mo, 6mo–12mo) for native **(i)** and IgA+ **(j)** fractions, compared against within-subject distances. Individual specificity in the IgA+ fraction was driven primarily by the 12-month timepoint (within-subject vs between-subject 12mo–12mo: *p* = 0.029) rather than the 6-month timepoint (*p* = 0.270). **k–n**, Paired PCoA trajectories connecting longitudinal samples from the same individual, plotted in the same ordination space as panels **(a)** and **(b)**. Lines connect timepoints within subjects and illustrate within-subject temporal change. Adults (**k**, native; **l**, IgA+) and infants (**m**, native; **n**, IgA+). Each color represents one subject. Violin plots show the data distribution; boxplots show the median (centre line), interquartile range (box bounds) and whiskers extending to 1.5× the interquartile range. \**p* < 0.05, \*\**p* < 0.01, \*\*\**p* < 0.001; ns, not significant.

**Figure 3.**
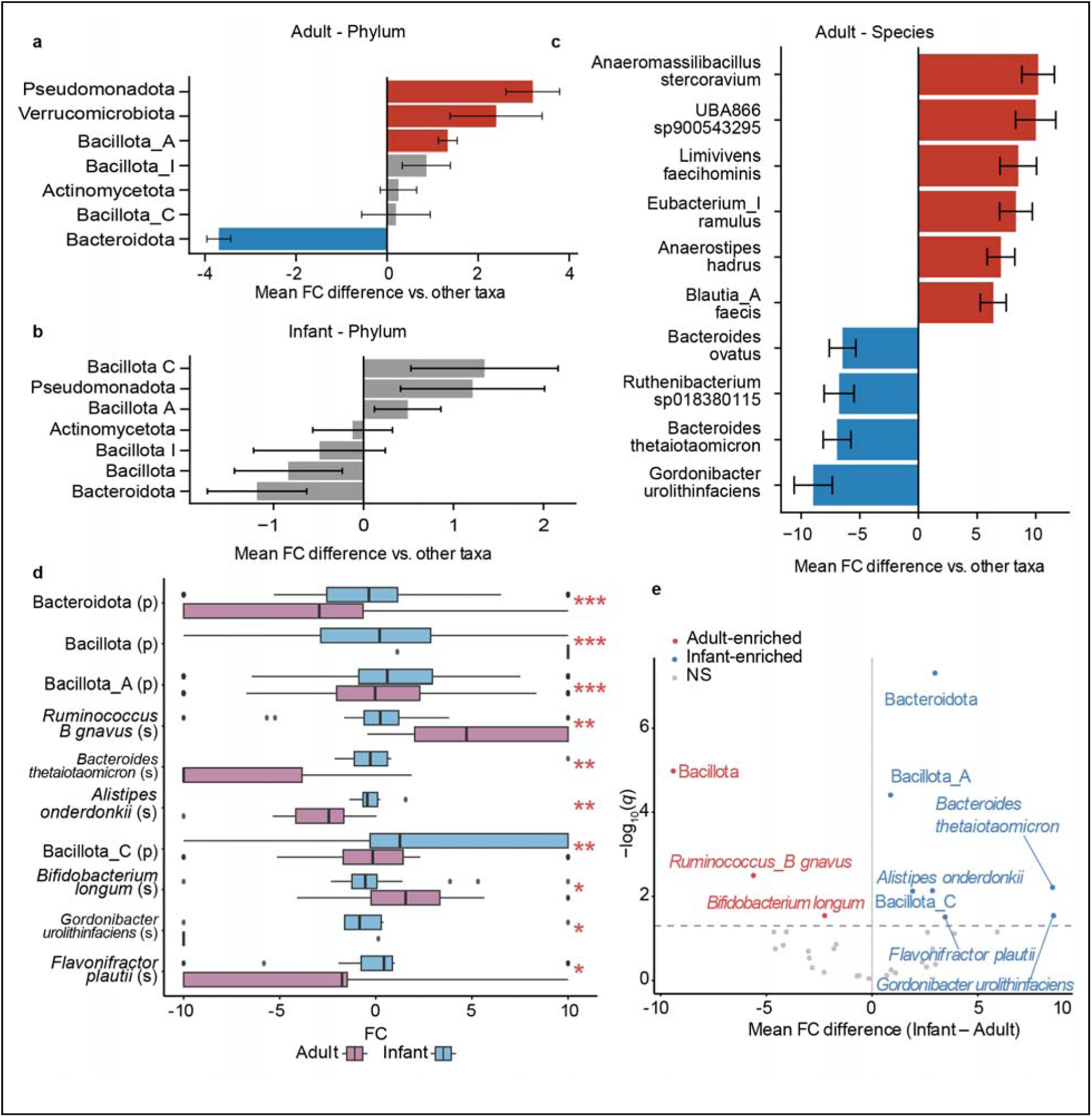
IgA targeting is conserved at the phylum level but diffuse at the species level in infants. **a,b,** Phylum-level taxa with significant differences in fold-change (FC) values relative to all other taxa within adults (**a**) and infants (**b**). Bars show mean FC difference; red indicates higher IgA targeting (positive FC) relative to other taxa, blue indicates lower IgA targeting (negative FC). Linear mixed-effects model (FC ∼ taxon identity + (1|subject); lmerTest) with Benjamini-Hochberg correction (*q* < 0.05) **c**, Species-level taxa with significantly different FC values in adults, colored as in (a,b). No individual species reached significance in infants. **d**, FC distributions for the 10 taxa with significantly different IgA targeting between infants and adults. Wilcoxon rank-sum test, Benjamini-Hochberg corrected. (s), species level; (p), phylum level. **e**, Mean FC difference (infant minus adult) versus significance (-log₁₀ q-value) for all taxa with sufficient observations in both age groups. Blue, significantly higher FC in infants; red, significantly higher FC in adults; grey, not significant. Horizontal dashed line indicates *q* = 0.05. Boxplots show the median (centre line), interquartile range (box bounds) and whiskers extending to 1.5× the interquartile range. \**q* < 0.05, \*\**q* < 0.01, \*\*\**q* < 0.001.

**Figure 4.**
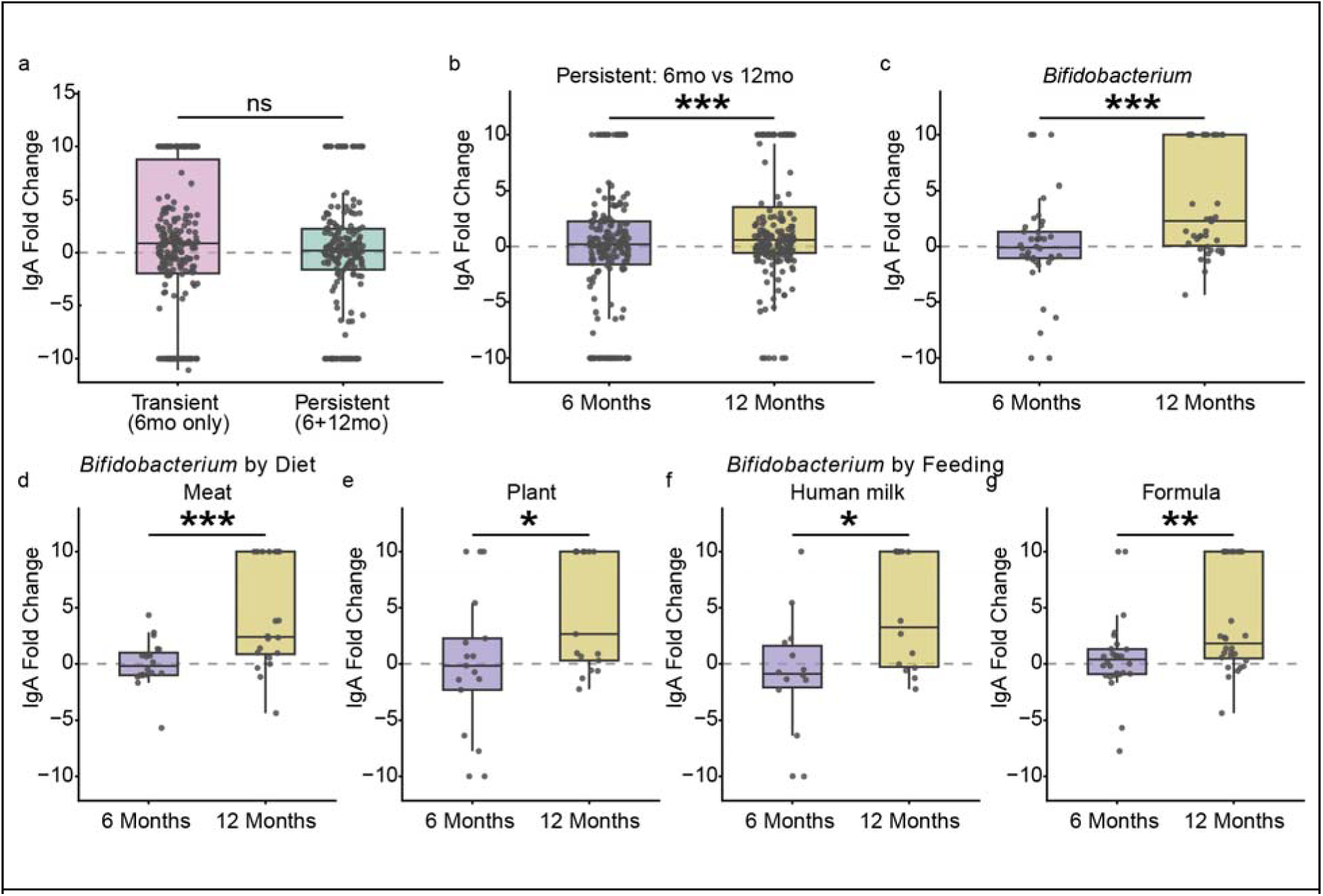
Persistent colonizers become progressively targeted by IgA. **a**, IgA fold-change at 6 months for transient taxa (detected at 6 months only) versus persistent taxa (detected at both 6 and 12 months). **b,** IgA fold-change of persistent taxa compared between 6 and 12 months. **c**, IgA fold-change of persistent Bifidobacterium species compared between 6 and 12 months. **d,e**, Persistent Bifidobacterium IgA fold-change at 6 versus 12 months stratified by dietary group: Meat **(d)** and Plant **(e)**. f,g, Persistent Bifidobacterium IgA fold-change at 6 versus 12 months stratified by feeding mode: human milk **(f)** and formula **(g)**. Linear mixed-effects model with subject as random intercept. \**p* < 0.05, \*\**p* < 0.01, \*\*\**p* < 0.001; ns, not significant. Boxplots show the median (centre line), interquartile range (box bounds) and whiskers extending to 1.5× the interquartile range. Each point represents one species-level observation.

Infant fecal samples were collected at home by caregivers, stored in a home freezer (−20°C) for up to one day, and then transported to the University of Colorado Anschutz, where they were aliquoted and stored at −80°C.

### 2. Metagenomic immunoglobulin sequencing (MIg-seq) for infant samples

The MIG-Seq protocol was adapted and optimized from previous methods to suit low-biomass infant fecal samples ^17, 16^.

#### Buffer Preparation

Staining Buffer: phosphate-buffered saline (PBS) supplemented with 1% bovine serum albumin (BSA; Sigma-Aldrich, #A7888). Used for all antibody dilutions, washes, and cell resuspensions. Blocking Buffer: Staining Buffer supplemented with 20% Normal Mouse Serum (Jackson ImmunoResearch, #015-000-120). Used exclusively during the pre-antibody blocking step to saturate non-specific binding sites on bacterial surfaces before anti-IgA-phycoerythrin (PE) incubation. Separation Buffer: PBS supplemented with 0.5% BSA and 0.5 mM ethylenediaminetetraacetic acid (EDTA). Used exclusively during magnetic bead washing steps. EDTA chelates divalent cations to inhibit deoxyribonuclease (DNase) activity and prevent bacterial aggregation during magnetic separation.

#### Sample Preparation and Fractionation

Approximately 200 mg of fecal material (acceptable range: 100-300 mg) was homogenized in 1 mL of ice-cold PBS and centrifuged at 500 × g for 15 minutes to pellet large debris. The supernatant (∼1 mL) was carefully collected and passed through a 70-μm cell strainer (Corning, #431751). From the filtered suspension, three aliquots were immediately reserved: (1) a 90 μL aliquot stored on ice as the ‘native’ fraction for total-community metagenomic sequencing; (2) a 5 μL aliquot combined with 195 μL Staining Buffer for bacterial quantification by flow cytometry; and (3) a 5 μL aliquot stained with 1× SYBR Green I (Invitrogen, #S7563) in 200 μL Staining Buffer to quantify background PE signal. The remaining ∼900 μL was retained for IgA labeling and magnetic enrichment.

#### IgA Labeling and Magnetic Enrichment

The remaining ∼900 μL of filtered suspension was divided equally between two wells of a deep-well 96-well plate (∼450 μL each). These duplicate wells were processed in parallel to maximize final biomass and combined at the end of the protocol. Wells were centrifuged at 3,000 × g for 9 minutes to pellet bacterial cells and supernatants were discarded.

Bacterial pellets were resuspended in 100 μL of Blocking Buffer and incubated for 20 minutes on ice. Following blocking, wells were washed with 1 mL of cold Staining Buffer and centrifuged at 3,000 × g for 9 minutes.

Pellets were then resuspended in 100 μL of anti-human IgA-PE antibody (clone IS11-8E10; Miltenyi Biotec, #130-113-476) diluted 1:10 in Staining Buffer, and incubated for 40-50 minutes on ice in the dark.

Following antibody incubation, samples were washed with 1 mL cold Staining Buffer and centrifuged at 3,000 × g for 9 minutes. Pellets were resuspended in 200 μL Staining Buffer. A 5 μL aliquot was taken and stained with 1× SYBR Green I in 200 μL Staining Buffer for flow cytometry analysis to quantify the baseline IgA-binding percentage in the unenriched (pre-sort) sample.

The remaining ∼195 μL was incubated with 20 μL of MojoSort Mouse Anti-PE Nanobeads (BioLegend, #480080) for 30 minutes at room temperature. MojoSort Mouse Anti-PE Nanobeads were selected following comparative testing against Miltenyi Anti-PE MicroBeads UltraPure (#130-105-639) and Invitrogen AccuCheck Counting Beads, as they demonstrated superior capture efficiency for low-biomass infant samples.

Samples were then placed on the Invitrogen Magnetic-Ring Stand (Invitrogen, #AM10050) for 5 minutes. The unbound supernatant was discarded, and the magnetically captured fraction was washed by removing the plate from the magnet, resuspending in 200 μL Separation Buffer, and returning to the magnet for 5 minutes. This wash cycle was repeated for a total of three magnetic enrichments and washes.

After the final wash, the duplicate wells for each sample were combined, yielding a final volume of ∼400 μL designated as the ‘IgA-positive’ fraction. A 5 μL aliquot was taken and stained with 1× SYBR Green I in 200 μL Staining Buffer for flow cytometry validation. The remaining ∼395 μL was reserved for DNA extraction and metagenomic sequencing.

#### Flow Cytometry Analysis

All flow cytometry analyses were performed using the MACSQuant VYB (Miltenyi Biotec). Four controls and samples were prepared for each specimen, all stained with 1× SYBR Green I (Invitrogen, #S7563) for 5 minutes at room temperature to identify bacterial cells by DNA staining: (1) an unstained control (native sample without SYBR Green or anti-IgA-PE), used to set background fluorescence thresholds; (2) a SYBR-only control (native sample stained with SYBR Green only), used to define the bacterial gate and establish background signal in the PE channel; (3) a native IgA-stained sample (unenriched native sample stained with both SYBR Green and anti-IgA-PE), used to quantify the baseline percentage of IgA-coated bacteria in the total community; and (4) an enriched IgA-positive sample (the final magnetically enriched fraction stained with SYBR Green), used to confirm enrichment efficiency.

Data were analyzed using FlowJo software (BD Life Sciences). The bacterial gate was established using side scatter (SSC) versus SYBR Green fluorescence intensity. The net percentage of IgA-coated bacteria was calculated by subtracting the background PE signal in the SYBR-only control from the PE-positive percentage in the native IgA-stained sample.

#### Metagenomic sequencing

DNA was extracted from the reserved native and IgA-positive fractions using the ZymoBIOMICS DNA Miniprep Kit (Zymo Research, #D4300) following the manufacturer’s protocol for low biomass samples. DNA concentration was quantified using a Qubit 2.0 Fluorometer (Invitrogen, #Q32866).

To select an optimal sequencing provider for our low-biomass samples, an initial validation was performed by submitting a subset of extracted DNA to three centers: Duke Sequencing and Genomic Technologies, Zymo Research Corporation, and Novogene Corporation. Based on a comparative analysis of data quality and performance, Duke Sequencing and Genomic Technologies was selected for sequencing the full cohort. All samples were subsequently sent to Duke for library preparation and shotgun metagenomic sequencing on an Illumina NovaSeq 6000 platform, generating 2 x 150 bp paired-end reads.

### 3. Bioinformatic Analysis

All bioinformatic analyses were conducted using a pipeline from the OlmLab/bioinformatics_pipelines Nextflow ^20^ repository (https://github.com/OlmLab/bioinformatics_pipelines). Raw paired-end reads from both ‘Pre-sorted’ (native) and ‘Post-sorted’ (IgA-positive) fractions were first processed using the roadmap_4 workflow for quality trimming, adapter removal (fastp) ^21^, and human host read decontamination (Bowtie2) ^22^. High-quality, non-host reads were then processed with the roadmap_6 workflow to generate community profiles. Specifically, species-level relative abundances for all samples were estimated using Sylph ^23^ against the Genome Taxonomy Database (GTDB, r220) ^24^.

### 4. Data analysis and statistical methods Quantification of IgA Targeting

IgA targeting strength for each microbial taxon in each sample was quantified using a log₂ fold-change (FC) score computed from paired native and IgA-positive sequencing data from the same fecal sample:

FC = log₂(IgA⁺ relative abundance / native relative abundance)

Taxa absent from both fractions were excluded (FC = NA). Taxa detected only in the IgA-positive fraction were assigned FC = +10, and taxa detected only in the native fraction were assigned FC = −10.

Flow cytometry data were used to validate enrichment efficiency at each processing step rather than as a primary quantitative input.

#### Beta-diversity and Community Structure Analysis

Samples collected at 5 months and 6 months of age were pooled as a single pre-intervention timepoint (“6 months”) for all analyses, as both preceded the dietary intervention. All adult comparison data were derived from the published MIg-Seq dataset ^17^. Beta-diversity was assessed using both unweighted and weighted UniFrac ^25^distances, computed using the UniFrac function in the phyloseq ^26^ R package (normalized = TRUE). Phylogenetic distances were calculated using a reference phylogenetic tree derived from the GTDB r220 bac120 marker gene set. Principal coordinates analysis (PCoA) was performed on the resulting distance matrices using cmdscale (k = 2, eig = TRUE). The effect of age group (6 months, 12 months, adult) and, separately, dietary intervention on community composition was tested using PERMANOVA ^27^ (adonis2, vegan package; 999 permutations). For comparisons between the 6- and 12-month infant timepoints, permutations were restricted within subjects using the strata argument to account for the repeated-measures design.

To assess individual specificity, we tested whether samples from the same individual were more similar to each other than to samples from different individuals. For each age group (adults and infants) and each fraction (native and IgA-positive), within-subject pairwise UniFrac distances (i.e., the distance between the two longitudinal samples from the same individual) were compared to between-subject distances (i.e., distances between samples from different individuals) using two-sided Wilcoxon rank-sum tests. To assess temporal stability across life stages, within-subject distances were compared between adults and infants using a two-sided Wilcoxon rank-sum test; larger within-subject distances indicate greater temporal change in community composition. All adult comparison data were derived from the published MIg-Seq dataset ^17^. Taxonomic assignments from that dataset (GTDB r202) were mapped to GTDB r220 nomenclature to ensure comparability with the infant samples.

#### Taxon-level IgA Targeting Comparisons

All taxon-level statistical analyses were performed on species- and phylum-level assignments derived from GTDB r220 taxonomy. IgA targeting strength (FC) was calculated per taxon per sample, such that each infant at each timepoint contributed one FC value for every detected taxon (see Quantification of IgA Targeting). Three sets of comparisons were conducted.

First, to identify taxa with unusually high or low IgA targeting within each age group (infants or adults), the FC values of each taxon were compared against the pooled FC values of all other taxa within the same age group. We note that these comparisons include multiple FC observations per subject (one per detected taxon at each level), which introduces within-subject correlation and violates the independence assumption of standard rank-sum tests. We therefore used linear mixed-effects models (LMM) as the primary analysis for these within-age comparisons, with subject as a random intercept to account for the repeated-measures structure: FC ∼ taxon_indicator + (1|subject_id), fitted using the lme4 and lmerTest packages in R. The taxon_indicator coefficient estimates the mean FC difference between the target taxon and all other taxa, with Satterthwaite-approximated degrees of freedom used for inference.

Second, to identify taxa with divergent IgA targeting between life stages, FC values for each taxon were compared between infants and adults using two-sided Wilcoxon rank-sum tests.

Minimum observation thresholds were set according to the sample size available for each comparison: at least 20 total observations and 3 subjects per taxon for phylum-level within-age LMM comparisons; at least 10 total observations and 3 subjects per taxon for species-level within-age LMM comparisons. P-values were corrected for multiple comparisons using the Benjamini–Hochberg procedure, applied separately within each comparison set. Taxa with q < 0.05 were considered statistically significant.

#### Persistence Analysis

To assess how IgA targeting relates to microbial colonization dynamics, we performed a persistence analysis across all 16 infants with paired samples at 6 and 12 months. A species was considered detected in a given sample if it had a non-zero relative abundance in either the native or IgA-positive fraction. For each infant, species detected at 6 months were classified as persistent (also detected at 12 months in the same subject) or transient (detected only at 6 months).

IgA fold-change (FC) values at 6 months were compared between transient and persistent taxa using linear mixed-effects models (LMM) with subject as a random intercept (FC ∼ persistence + (1|subject_id)), with Persistent as the reference level, fitted using the lme4 and lmerTest packages with Satterthwaite-approximated degrees of freedom. To assess longitudinal changes in IgA targeting of persistent colonizers, FC values for persistent species were compared between 6 and 12 months using LMM (FC ∼ timepoint + (1|subject_id)), with 6mo as the reference level. These comparisons were further stratified by dietary group (Meat vs Plant) and feeding mode (human milk vs formula) using the same LMM framework. Since only a small number of statistical tests were performed in the persistence analysis (N = 7), nominal p-values are reported without multiple-testing correction.

### Diet and Covariate Analyses

#### Baseline covariate and arm comparability checks

To confirm that IgA targeting did not differ systematically by demographic characteristics prior to dietary intervention, phylum-level IgA FC at enrollment (5–6 months) was compared across clinical covariates (sex, delivery mode, childcare attendance) using two-sided Wilcoxon rank-sum tests on per-subject mean phylum-level FC, with BH correction applied within each covariate across phyla. Infants with mixed feeding (feeding_method = 3) were excluded from the baseline feeding mode comparison, which was restricted to exclusively human milk-fed and exclusively formula-fed infants. Randomization balance across the four dietary arms was assessed using Kruskal-Wallis tests on per-subject mean phylum-level FC, with BH correction across phyla. At 12 months, associations between phylum-level IgA FC and clinical covariates (sex, delivery mode, feeding mode, childcare attendance) were assessed using LMM (FC ∼ covariate + (1|subject_id)), fitted separately for each covariate, with BH correction across phyla within each covariate. The overall level of IgA activity at 12 months was compared between Plant and Meat arms using a two-sided Wilcoxon rank-sum test on per-subject median FC across all detected taxa, yielding one value per subject and avoiding independence violations.

#### Dietary nutrient-IgA targeting correlations

Three-day weighed diet records were analyzed using the Nutrition Data System for Research (NDSR, University of Minnesota), averaged to daily nutrient intakes per subject by summing all recorded intakes and dividing by the number of available recording days; subjects with no dietary records were excluded (n = 14 retained). Two subjects with fewer than three recording days (one with 1 day, one with 2 days) were included using the same averaging approach, with the caveat that their daily intake estimates are based on fewer observations and may be less representative of habitual intake. Nutrient columns were subjected to a quality filter requiring non-zero variance, ≥8 non-missing values, and <80% zero values across subjects, retaining 153 nutrients for analysis. Per-subject mean IgA FC was calculated for each species detected in ≥8 of the 14 subjects with dietary data at 12 months. Spearman’s rank correlation (ρ) was computed for each nutrient × species pair; BH correction was applied within each nutrient across all tested species, and associations with *q* < 0.10 were considered noteworthy.

### Comparison of nutrient associations with IgA targeting and native relative abundance

To assess whether nutrient–IgA FC associations reflected IgA targeting specificity rather than covariation with microbiome composition, a parallel Spearman correlation analysis was performed using native relative abundance from the unsorted metagenomic fraction as the outcome variable in place of IgA FC. Per-subject mean relative abundance was calculated for each species detected in ≥8 subjects with dietary data at 12 months, consistent with the IgA FC analysis. The same set of 153 quality-filtered nutrients and BH correction procedure (applied within each nutrient across all tested species) were used, yielding 1,683 total tests. To compare results across the two analyses, all nutrient–species pairs were categorized according to significance (*q* < 0.10) in each analysis: significant in IgA FC only, in relative abundance only, in both, or in neither. For all pairs significant in at least one analysis, the direction of association (concordant vs. opposing sign of Spearman’s ρ) was recorded.

### Bifidobacterium-dietary fiber and nutrient associations

The association between total dietary fiber intake and Bifidobacterium IgA targeting was examined at two levels of resolution. At the subject level, per-subject mean Bifidobacterium FC was calculated by averaging FC values across all detected Bifidobacterium species within each subject, and correlated with daily total dietary fiber intake using Spearman’s ρ (n = 14). At the species level, all Bifidobacterium species-level FC observations were analyzed using LMM (FC ∼ Total_Dietary_Fiber_g + (1|subject_id)) to account for within-subject correlation across multiple detected species. A complementary systematic screen was conducted by correlating the same per-subject mean genus-level Bifidobacterium FC against all 153 quality-filtered nutrients using Spearman’s ρ, with BH correction applied across nutrients; this genus-level approach averaged across all detected Bifidobacterium species per subject without applying a per-species detection threshold.

All statistical analyses were conducted in R (v 4.5.2)

## Results

### Performing MIg-Seq on Infant Fecal Samples

MIg-Seq has previously been used to characterize IgA-microbiome interactions in adults^17^, but has not been previously applied to infant fecal samples. We made several modifications to adapt the protocol to perform on lower-biomass infant fecal samples. Key modifications included 1) a dedicated pre-incubation blocking step with normal mouse serum prior to antibody labeling, replacing the co-incubation approach used in adult protocols, 2) an increased anti-IgA-PE antibody concentration (1:10 versus 1:30) to ensure sufficient signal strength, 3) the incorporation of EDTA into the magnetic separation buffer to minimize bacterial aggregation during washes, and 4) changing the fluorophore from APC to PE (**Fig. 1a**). See Methods for details.

We next applied the updated Metagenomic Immunoglobulin Sequencing (MIg-Seq) protocol to 32 paired longitudinal fecal samples from 16 infants enrolled in a randomized controlled trial (RCT) of complementary feeding strategies **(Fig. 1b)** ^28^. Complementary feeding is the period from around 6-12 months of age where solid foods are introduced alongside the infant’s primary diet of breast-milk / formula, and this RCT is the first ever performed to directly evaluate the impact of meat-, vegetable-, or dairy-enriched complementary feeding on infant health outcomes when compared to a reference (no intervention) group. The intervention supplied protein-rich foods from arm-specific sources (meat, dairy, or plant) targeting a specific protein intake (∼84 kcal/d out of an estimated ∼650 kcal/d total complementary calories), while remaining dietary choices were at caregiver discretion.

Infants were block randomized into dietary groups at 5 or 6 months of age (prior to the start of complementary feeding) and continued the intervention until 12 months of age. For this study, we focused on infants receiving meat-based (n = 6) and plant-based (n = 8) diets, and included one infant each from the dairy-based and reference groups to maximize sample size for non-diet-specific analyses. Among the 15 infants with available clinical metadata, the cohort was predominantly female (71%) and delivered vaginally (57%). Seven infants were exclusively human milk-fed and 9 were exclusively formula-fed at the time of MIg-Seq sampling; the distribution across diet and feeding mode groups is reported in **Table 1**. Sample sizes for covariate and dietary analyses varied by data availability and are detailed in the Methods (n = 15 for covariate analyses; n = 14 for diet–IgA correlation analyses).

Flow cytometry confirmed the specificity of the anti-IgA-PE antibody labeling, with unstained controls showing negligible background signal across all samples **(Fig. 1c,d)**. The proportion of IgA-coated bacteria in the native community varied substantially across individuals and timepoints **(Fig. 1d)**, as has been previously seen in adult fecal samples ^17^. To detect low-abundance microbes, we also performed relatively deep shotgun sequencing on both the native and IgA-enriched microbial fractions (Sequencing depth = 8.0 ± 4.0 Giga base-pairs). Taken together, this dataset represents a one-of-a-kind opportunity to study the impact of early diet and human milk feeding on the infant mucosal immune system.

### Infant IgA targeting exhibits individual specificity

To establish baseline features of infant IgA–microbiome interactions, we compared the 32 infant samples to 30 longitudinal adult fecal samples previously characterized by MIg-Seq^17^. Adult samples were collected before and after an 8-week dietary intervention that did not significantly alter IgA targeting.

PCoA of unweighted UniFrac distances revealed clear separation between infant and adult samples in both the native and IgA+ fractions **(Fig. 2a,b)**, confirmed by pairwise PERMANOVA (native: R² = 0.329, *p* = 0.001; IgA+: R² = 0.285, *p* = 0.001; **Supplementary Table S1**). Within infants, 6-month and 12-month samples largely overlapped in ordination space; timepoint explained a modest but non-significant proportion of variance in the native fraction (R² = 0.047, *p* = 0.075) and a small but significant proportion in the IgA+ fraction (R² = 0.056, *p* = 0.004). Weighted UniFrac showed consistent results (IgA+ *p* = 0.028; native *p* = 0.153; **Supplementary Table S1**).

In adults, within-subject UniFrac distances were significantly smaller than between-subject distances in both the native and IgA+ fractions (native: *p* = 1.2×10⁻; IgA+: *p* = 2.3×10⁻; **Fig. 2c,d**), confirming stable, individual-specific community profiles. In infants, within-subject distances were significantly smaller than between-subject distances in the IgA+ fraction (*p* = 0.040; **Fig. 2f**) but not in the native fraction (*p* = 0.336; **Fig. 2e**), indicating that IgA targeting is more stable over time than overall microbiome composition. To determine when this specificity emerges, we stratified between-subject distances by timepoint pair (**Fig. 2i, j**). In the IgA+ fraction, individual specificity was significant at 12 months (within vs between 12mo–12mo: *p* = 0.029) but not at 6 months (within vs between 6mo–6mo: *p* = 0.270). Comparing within-subject distances to cross-timepoint between-subject distances (one infant’s 6mo vs another infant’s 12mo sample) also reached significance (*p* = 0.020), indicating that self-similarity in IgA targeting persists across the 6-to-12-month interval. Weighted UniFrac results were consistent (**Supplementary Table S1**). Paired longitudinal trajectories are shown in **Fig. 2k–n**. Within-subject temporal distances were greater in infants than in adults for both fractions (native: *p* = 3.9×10⁻³; IgA+: *p* = 7.8×10⁻; **Fig. 2g,h**), but this comparison should be interpreted with caution given the longer longitudinal interval in infants (∼6 months) compared to adults (8 weeks).

### Infant IgA targeting generally mirrors adult binding

To characterize taxon-level IgA targeting, we computed a fold-change (FC) score for each taxon reflecting its relative enrichment in the IgA+ versus native fraction (see Methods). Linear mixed-effects models (LMM) with a subject-level random intercept were used to identify taxa with FC values significantly deviating from all other taxa within each age group.

At the phylum level, the directionality of IgA targeting was broadly conserved between infants and adults **(Fig. 3a, b)**: Bacteroidota was consistently depleted in the IgA+ fraction, while Bacillota_A and Pseudomonadota were enriched. These trends reached statistical significance only in adults (Bacteroidota: *q* = 4.2×10⁻ ³; Bacillota_A: *q* = 4.3×10⁻¹; Pseudomonadota: *q* = 1.3×10⁻; Verrucomicrobiota: *q* = 0.031); no phylum reached significance in infants after BH correction. At the species level, 55 taxa showed significantly high or low FC values in adults **(Fig. 3c)**, while no individual species reached significance in infants, despite comparable sample sizes (n = 32 infant, n = 30 adult) and a longer longitudinal interval (6 months versus 8 weeks).

Direct comparison of FC values between age groups identified 10 taxa with significantly different IgA targeting between infants and adults **(Fig. 3d, e)**. The largest phylum-level divergences were observed for Bacteroidota, Bacillota, and Bacillota_A (all *q* < 0.001). At the species level, *Ruminococcus_B gnavus* (*q* = 3.1×10⁻³), *Bacteroides thetaiotaomicron* (*q* = 6.1×10⁻³), and *Bifidobacterium longum* (*q* = 0.029) showed the greatest life-stage differences. No individual taxon showed significantly different FC between 6 and 12 months within infants, despite the significant community-level shift in the IgA+ fraction detected by PERMANOVA.

### Progressive IgA targeting of persistent colonizers is driven by Bifidobacterium

To characterize how IgA targeting evolves with microbial colonization dynamics, we classified taxa as transient (detected only at 6 months) or persistent (detected at both timepoints) and compared IgA FC between groups using LMM with the subject as a random intercept. Transient taxa showed higher FC at 6 months than persistent taxa (median FC: 0.88 vs 0.20), but this difference was not significant (*p* = 0.473; **Fig. 4a)**. Among persistent taxa, IgA FC increased significantly from 6 to 12 months (median FC: 0.20 to 0.59; *p* < 0.001; **Fig. 4b**), indicating progressive strengthening of IgA targeting toward stable community members.

This effect was most pronounced for *Bifidobacterium*, which showed a highly significant increase in IgA FC from 6 to 12 months (median FC: −0.09 to 2.29; *p* < 0.001; **Fig. 4c**). The increase was consistent across dietary groups (Meat: *p* < 0.001; Plant: *p* = 0.025; **Fig. 4d,e**) and feeding modes (human milk: *p* = 0.011; formula: *p* = 0.004; **Fig. 4f, g**).

### Complementary feeding impacts IgA targeting

We first tested whether demographic variation and/or clinical covariates impacted IgA targeting. At both the 5 months and 12 months timepoints, we found no significant association between IgA binding of any bacterial phylum and sex, delivery mode, or childcare attendance (Wilcoxon rank-sum test per phylum; all BH-corrected *q* > 0.50; **Supplementary Table S4**). We next assessed the impact of feeding mode (breastmilk versus formula) on community composition and IgA targeting. At 6 months of life neither the native nor IgA+ fractions showed significant separation by feeding mode across any distance metric (PERMANOVA, all p > 0.05; **Supplementary Fig. S1a-d**), though the small sample size (n = 7 human milk-fed, n = 9 formula-fed) limits power to detect modest effects. The overall proportion of IgA-coated bacteria trended higher in breastmilk-fed infants (32.9 ± 11.0%) compared to formula-fed infants (25.7 ± 9.8%), but this difference was not significant. At the taxonomic level, no species showed significantly different FC values after FDR correction **(Supplementary Table S6)**.

Similarly, after 6 months of dietary intervention, overall community composition remained indistinguishable between dietary groups across all distance metrics and fractions (PERMANOVA, all p > 0.05; **Supplementary Fig. S1 e-l**). Using a FDR correction threshold of 0.1, no significant differences in IgA binding were linked to the dietary intervention **(Supplementary Table S7)**. The overall magnitude of IgA targeting did not differ between the Plant and Meat arms at 12 months (median FC: 0.60 vs. 0.86; Wilcoxon W = 20, p = 0.648). Kruskal-Wallis testing confirmed comparable IgA targeting across all four dietary arms at baseline and 12 months (all *p* > 0.43, *q* > 0.75; **Supplementary Table S5**).

We next used continuous nutrient intake data from available 3-day dietary records (n = 14; collected prior to the 12-month sampling timepoint) to identify specific dietary drivers of IgA targeting. This approach was motivated by the fact that the plant vs. meat arm assignment controlled only the source of protein-rich complementary foods, which contributed a minority of total complementary calories (∼25%), while the remaining complementary diet was uncontrolled and similar across arms. We correlated per-subject daily nutrient intakes with species-level IgA FC using Spearman’s ρ, with BH correction applied within each nutrient across all species (**Fig. 5A**; **Supplementary Table S8**). Among all nutrient–species pairs tested, 50 associations met *q* < 0.10 (Table 2; Supplementary Table SX for complete results). The strongest association was between cholesterol intake and IgA targeting of *Bifidobacterium longum* (ρ = 0.909, *q* = 0.001; **Fig. 5B)**; cholesterol was elevated in the Meat arm relative to the Plant arm (Wilcoxon *p* = 0.041). Additional animal-derived nutrients like animal protein, EPA (20:5 n-3), and both forms of conjugated linoleic acid, showed concordant positive associations with *B. longum* FC (all *q* < 0.05; **Table 2**).

**Figure 5.**
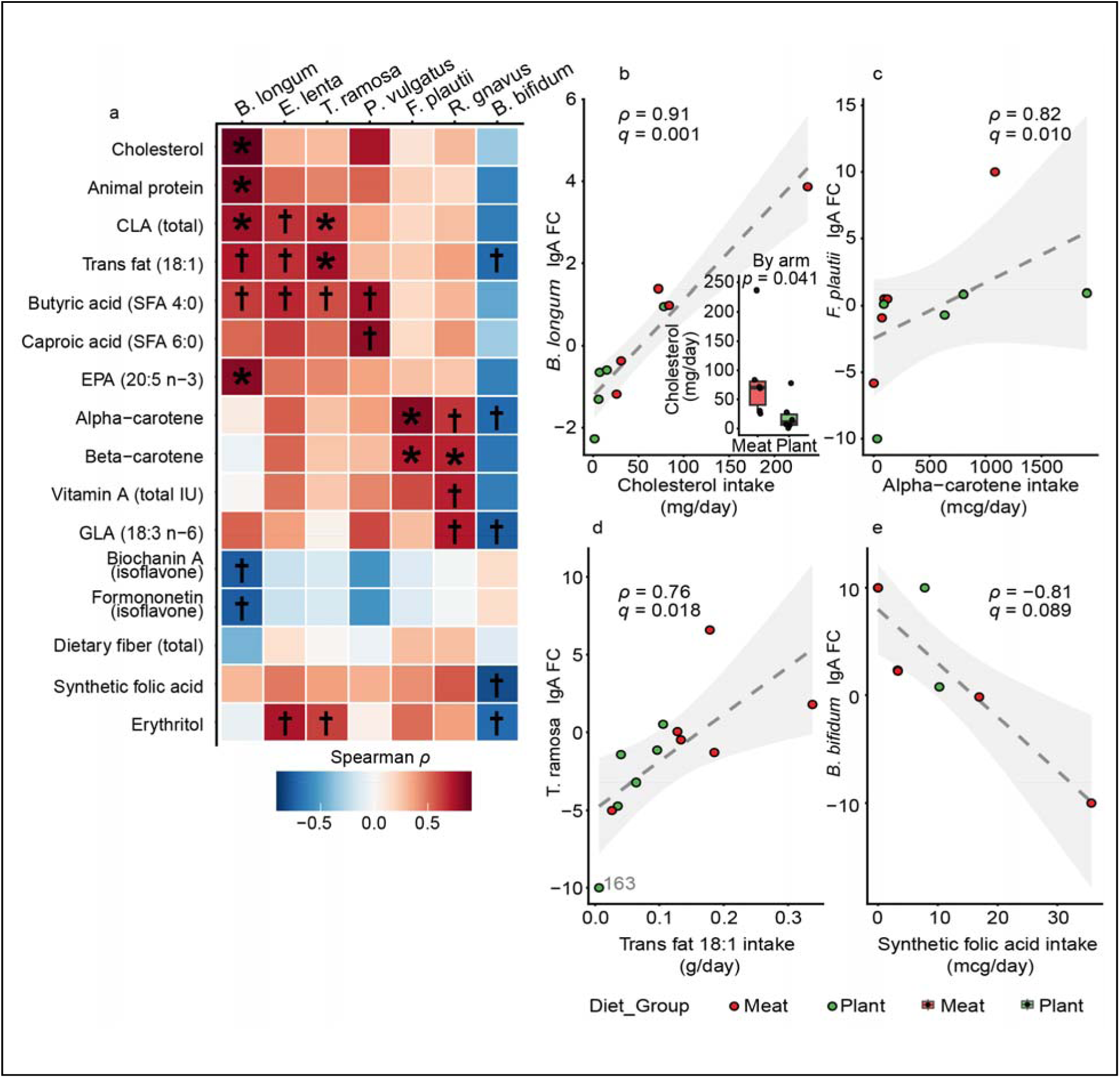
Complementary diet reshapes IgA targets through specific nutrient–taxon associations. **a,** Heatmap of Spearman correlations between dietary nutrient intake and species-level IgA FC at 12 months. Only nutrients with at least one association at *q* < 0.10 are shown. Color scale indicates Spearman ρ; \**q* < 0.05, †*q* < 0.10 (BH correction applied within each nutrient across all species; n per species reported in **Table 2**). **b**, Cholesterol intake versus *B. longum* IgA FC (ρ = 0.91, *q* = 0.001); inset shows cholesterol intake by dietary arm (Wilcoxon *p* = 0.041). **c**, Alpha-carotene intake versus *F. plautii* IgA FC (ρ = 0.82, *q* = 0.010). **d**, Trans fat 18:1 intake versus *T. ramosa* IgA FC (ρ = 0.76, *q* = 0.018). **e**, Synthetic folic acid intake versus *B. bifidum* IgA FC (ρ = −0.81, *q* = 0.089). Points colored by dietary group (red, Meat; green, Plant). Dashed line, linear fit; shading, 95% confidence interval.

**Table 2.** Strongest dietary nutrient predictor of IgA binding per species at 12 months.

| Nutrient | Species | N | $\rho$ | p-value | q-value |
| --- | --- | --- | --- | --- | --- |
| **Cholesterol** | B. longum | 11 | 0.909 | <0.001 | 0.001160 |
| **CLA (trans-10, cis-12)** | B. longum | 11 | 0.874 | <0.001 | 0.004760 |
| **Alpha-carotene** | F. plautii | 12 | 0.825 | <0.001 | 0.01050 |
| **Animal protein** | B. longum | 11 | 0.836 | 0.001330 | 0.01470 |
| **EPA (20:5 n-3)** | B. longum | 11 | 0.828 | 0.001640 | 0.01810 |
| **Trans fat (18:1)** | T. ramosa | 14 | 0.758 | 0.001670 | 0.01840 |
| **Beta-carotene** | F. plautii | 12 | 0.720 | 0.008240 | 0.04530 |
| **Beta-carotene** | R. gnavus | 13 | 0.703 | 0.007320 | 0.04530 |
| **CLA (18:2, total)** | B. longum | 11 | 0.770 | 0.005570 | 0.04570 |
| **CLA (18:2, total)** | T. ramosa | 14 | 0.673 | 0.008310 | 0.04570 |
| **Solid Fats** | B. longum | 11 | 0.780 | 0.004640 | 0.05110 |
| **GLA (18:3 n-6)** | R. gnavus | 13 | 0.729 | 0.004690 | 0.05160 |
| **CLA cis 9. trans 11** | B. longum | 11 | 0.733 | 0.01020 | 0.05610 |
| **CLA cis 9. trans 11** | T. ramosa | 14 | 0.667 | 0.009220 | 0.05610 |
| **Erythritol** | E. lenta | 12 | 0.746 | 0.005360 | 0.05900 |
| **Trans fat (18:1)** | B. longum | 11 | 0.727 | 0.01120 | 0.06160 |
| **Trans fat (18:1)** | E. lenta | 12 | 0.671 | 0.01680 | 0.06170 |
| **CLA (18:2, total)** | E. lenta | 12 | 0.662 | 0.01900 | 0.06970 |
| **Total Trans Fatty Acids TRANS** | T. ramosa | 14 | 0.688 | 0.006540 | 0.07190 |
| **PUFA 20.4 arachidonic acid. undifferentiated** | B. longum | 11 | 0.761 | 0.006550 | 0.07200 |

Among the nutrients associated with consumption of plants, alpha-carotene was the strongest plant-associated predictor of IgA targeting (ρ = 0.825 for Flavonifractor plautii, *q* = 0.011; **Fig. 5C**), with beta-carotene positively associated with IgA targeting of both *F. plautii* and *Ruminococcus gnavus* (both *q* = 0.045). Isoflavones were negatively associated with *B. longum* IgA FC (both ρ = −0.758, *q* = 0.075; Table 2). In contrast, total dietary fiber showed no association with *Bifidobacterium* IgA FC at the subject level (Spearman ρ = −0.174, *p* = 0.553, n = 14). An LMM incorporating all species-level observations with subject as a random effect yielded concordant null results (β = −0.140, 95% CI: −0.574 to 0.293, *p* = 0.503), ruling out confounding from unequal species representation. A systematic screen of all 153 quality-filtered nutrients identified carotenoids as the strongest negative correlates of genus-level *Bifidobacterium* IgA FC (alpha-carotene: ρ = −0.684, *p* = 0.007; beta-carotene: ρ = −0.640, *p* = 0.014), though none survived BH correction (all *q* > 0.50; **Supplementary Table S11**).

To assess whether the nutrient–IgA FC associations reflect genuine immune targeting specificity rather than passive covariation with microbiome composition, we performed a parallel Spearman correlation analysis substituting species-level IgA FC with native relative abundance from the unsorted metagenomic fraction, applying the same nutrient set and BH correction procedure. This analysis yielded only 9 significant associations (q < 0.10; **Supplementary Table S9**), substantially fewer than the 50 identified in the IgA FC analysis. Of the 50 FC-significant associations, 46 showed no corresponding relative abundance association (q ≥ 0.10), and the remaining 4 were significant in both analyses but with opposing direction in each case (**Supplementary Table S10**). Erythritol intake was positively associated with IgA targeting of *E. lenta* (ρ = 0.746, q = 0.059), *T. ramosa* (ρ = 0.628, q = 0.088), and *C. innocuum* (ρ = 0.638, q = 0.089), yet negatively associated with the relative abundance of each species (ρ = −0.711, q = 0.048; ρ = −0.654, q = 0.061; ρ = −0.590, q = 0.097, respectively) (**Supplementary Fig. S2)**. Similarly, solid fat intake was positively associated with *B. longum* IgA FC (ρ = 0.780, q = 0.051) and negatively associated with *B. longum* relative abundance (ρ = −0.703, q = 0.056). Five associations were significant (q < 0.10) only in the relative abundance analysis:medium-chain saturated fatty acids, including myristic acid (ρ = −0.717, q = 0.043), lauric acid (ρ = −0.713, q = 0.046), and caprylic acid (ρ = −0.656, q = 0.061), were each negatively associated with *B. longum* relative abundance, yet none showed a significant IgA FC association with *B. longum* (all q > 0.21). Taken together, these findings indicate that dietary nutrients reshape IgA targeting independently of, and in most cases inversely to, their effects on microbial colonization.

## Discussion

The MIg-Seq framework presented here extends species-level IgA profiling to the infant gut, a setting where low microbial biomass and low IgA-coating proportions have previously precluded metagenomic characterization of mucosal immune targeting. Applied to a randomized dietary intervention, our findings demonstrate that individual specificity in IgA targeting emerges progressively over the first year of life, that IgA targeting strengthens toward persistent colonizers in a manner that is most pronounced for *Bifidobacterium* and independent of diet or feeding mode, and that complementary feeding selectively reshapes IgA targets through specific nutrient–taxon associations without broadly altering overall IgA activity. Future species-resolution studies of IgA targeting across global infant cohorts, spanning healthy infants and those with immune-mediated disease, will be essential for understanding how early nutritional exposures shape the trajectory of mucosal immunity.

The broad phylum-level similarity between infant and adult IgA targeting suggests that certain broad microbial features, such as LPS architecture in *Pseudomonadota* or surface polysaccharide profiles in *Bacteroidota*, serve as consistent targets of mucosal immune recognition across the lifespan **(Fig. 3a,b)** ^29,5,30^. The near-absence of species-level targeting in infants, despite comparable sample sizes, likely reflects a lack of affinity-matured, T cell-dependent IgA. The apparent diffuseness of infant IgA targeting may represent an evolved permissiveness that enables a diverse and stable microbiome to become established before the system begins to sharpen its resolution. The three taxa with the most significantly divergent IgA targeting between infants and adults **(Fig. 3d,e)** are well characterized species: *B. longum* is a dominant human milk oligosaccharide (HMO) consumer in the human milk-fed infant gut ^31,32^, *Bacteroides thetaiotaomicron* is a versatile glycan forager with the capability to evade IgA recognition through phase variation of its surface capsular polysaccharides ^33^, and *Ruminococcus_B gnavus* is a mucin-degrading species with well-documented associations with aberrant IgA responses in adult inflammatory bowel disease ^15,34,35^. Whether the divergent IgA targeting of these species reflects their distinct ecological niches in the infant versus adult gut, differences in the strains that colonize each life stage, or simply the difference between new versus established colonization remains an open question.

The noted lack of within-subject relative to between-subject variation of microbiome composition **(Fig. 2e)** is consistent with the instability of the gut microbiota during the first year of life ^36–39^. The finding that the IgA-positive fraction exhibited significantly lower within-subject than between-subject distances **(Fig. 2f)** indicates that the host immune system confers a layer of individual-specific structure onto this turbulent community. Stratification by timepoint revealed that this specificity was detectable at 12 months (**Fig. 2j**, *p* = 0.029) but not at 6 months (*p* = 0.270), suggesting that individual specificity in IgA targeting emerges progressively over the first year rather than being present from the outset.This observation raises the question: is this emerging individual specificity in IgA targeting a property of the infant’s own emerging mucosal immune system, or is it largely inherited from maternal secretory IgA (sIgA) transferred via human milk? Maternal sIgA is shaped by decades of antigen exposure and germinal center reactions and represents a relatively stable, individual-specific antibody repertoire ^12,40,41^. Maternal sIgA has been hypothesized to act as a consistent immunological filter that trains the infant immune system to target specific microbial taxa ^42–44^. Alternatively, the specificity could reflect the infant’s own T cell-dependent IgA production, suggesting the neonatal mucosal immune system establishes targeted recognition earlier than previously appreciated ^42,45^. The progressive emergence of individual specificity between 6 and 12 months, coinciding with the period of active complementary feeding, is consistent with either mechanism, and distinguishing between them represents an important direction for future work.

The progressive strengthening of IgA targeting of persistent colonizers from 6 to 12 months, and the particularly pronounced increase for Bifidobacterium, constitutes a key finding that is robust across both dietary groups and both feeding modes **(Fig. 4)**. We hypothesize that stable colonization of these taxa results in the generation of antigen-specific B cell clones in gut-associated lymphoid tissue, progressively sharpening targeting toward established community members. This IgA binding trajectory of *Bifidobacterium* is especially interesting: we observe increased IgA targeting from 6 to 12 months in infants (**Fig. 5c**) and increased binding in adults relative to infants **(Fig. 3d)**. Given that *Bifidobacterium* is widely recognized as a health-associated early colonizer ^14,31,32,46^, we hypothesize that this strengthening IgA response is intended to promote *Bifidobacterium* colonization by anchoring to the mucus layer, as has been observed with *B. fragilis* in mouse models ^4^.

The substantially greater number of significant nutrient-IgA binding associations (50) compared to nutrient-relative abundance associations (9) underscores the added value of MIg-Seq over conventional microbiome taxonomy profiling for detecting diet-immune-microbiome interactions. This pattern mirrors findings from a previous study in adults, where IgA binding scores demonstrated far stronger correlations with dietary and immune variables than native relative abundance measures^17^. Together, these results suggest that IgA targeting captures a layer of host-microbiome interaction that is largely invisible to abundance-based approaches, and that dietary modulation of mucosal immunity may be underestimated by studies relying on microbiome composition alone.

A central finding of this study is that specific dietary nutrients, such as cholesterol and carotene, are significantly associated with the IgA targeting of specific bacterial taxa **(Fig. 5)**. The finding that *B. longum* binding is associated with animal-derived nutrient consumption, while *B. longum* abundance is anti-associated with animal-derived nutrient consumption, is particularly intriguing given that *B. longum* is a keystone member of the infant gut microbiome ^14,31,32,46^.One explanation is that meat-based complementary feeding alters the intestinal environment in ways that affect *B. longum* colonization dynamics or surface antigen presentation, secondarily driving IgA targeting; alternatively, cholesterol-associated metabolites may more directly influence mucosal B cell activity or IgA affinity maturation toward this taxon. The finding that *Thomasclavelia ramosa*, recently reclassified from *Erysipelatoclostridium ramosum* in GTDB r220 and identified as a target of secretory IgA-mediated immune exclusion in human milk ^43^, showed significant positive associations with animal intake is also particularly interesting (**Table 2**; **Fig. 5d**). The null result that total dietary fiber showed no detectable association with Bifidobacterium IgA FC at either the subject level (ρ = −0.174, *p* = 0.553) or in species-level mixed-effects models (β = −0.140, *p* = 0.503) is notable given the prominent hypothesis that fiber-derived short-chain fatty acids promote luminal IgA secretion and mucosal homeostasis ^47,48,49,50^. Potential explanations could include that fiber-derived SCFAs influence bulk IgA production without determining targeting specificity, or that this mechanism is not applicable in the context of the infant microbiome.

Future work should prioritize expanding sample size and longitudinal resolution, including infants from additional lifestyles, and exploring additional profiling techniques. Our core longitudinal cohort of 16 infants is sufficient to detect large-effect dietary responses, but has limited the power to detect subtle species-level associations or to fully account for the substantial inter-individual variability characteristic of the infant gut microbiome. Infant microbiomes are also known to widely vary in countries living different lifestyles ^51^, and understanding the impact of lifestyle on IgA binding would be especially interesting. Coupling MIg-Seq with paired human milk samples could also enable direct testing of whether the individual specificity observed in infant IgA-positive fractions is maternally derived, and mechanistic follow-up using germ-free mouse models colonized with defined infant microbial communities could establish causal relationships between specific dietary components, IgA secretion, and selective microbial targeting, as well as determine whether these interactions carry downstream phenotypic or metabolic consequences. Finally, while MIg-Seq applied to fecal samples captures the immunological landscape of the luminal compartment; the relationship between luminal IgA-microbiome interactions and those occurring at the mucosal surface, where the immunological stakes are highest, remains to be directly determined.

### Conclusions

Here we present the first species-resolution characterization of IgA–microbiome interactions during infancy, enabled by adapting Metagenomic Immunoglobulin Sequencing (MIg-Seq) for low-biomass infant fecal samples **(Fig. 1)**. Comparing infant profiles to a previously characterized adult cohort demonstrates that while phylum-level IgA targeting patterns are broadly conserved across life stages, the species-level selectivity observed in adults has not yet emerged in infants **(Fig. 3)**. We further show that the IgA-coated fraction of the microbiome is more individualized than the microbiome composition itself (**Fig. 2**). Strikingly, we also find that IgA targeting of persistent colonizers strengthens significantly from 6 to 12 months, with the most pronounced and consistent effect observed for *Bifidobacterium*, a finding robust across complementary feeding arms and human milk feeding status (**Fig. 4**). At the dietary level, complementary feeding selectively reshapes which taxa are targeted by IgA without altering overall IgA magnitude, and these effects are nutrient-specific: animal-derived nutrients, particularly cholesterol, are strongly associated with enhanced IgA targeting of *Bifidobacterium longum*, while plant-derived carotenoids are linked to increased targeting of *Flavonifractor plautii* and *Ruminococcus gnavus* (**Fig. 5**). Together, these findings establish a new immunological dimension through which to understand microbiome assembly and its responsiveness to early-life environmental exposures. These findings also lay the groundwork for developing IgA-based strategies, including monoclonal antibodies, vaccines, and modifiable lifestyle interventions, to therapeutically shape the infant microbiome and reduce the burden of immune disease in the industrialized world.

## Data availability

The data supporting the findings of this study are available within the paper and its Supplementary Information. Metagenomic reads are in the process of being uploaded to the NCBI Short Read Archive and the BioProject accession will be made publicly available.

## Code availability

nf-metgenomics-piplines: https://github.com/OlmLab/bioinformatics_pipelines

## Acknowledgements

We thank Sean P. Spencer for guidance on MIg-Seq protocol optimization. We thank the families who participated in the clinical trial and the clinical staff at the University of Colorado Anschutz Medical Campus for sample collection. We acknowledge the Department of Molecular, Cellular & Developmental Biology Flow Cytometry Facility and the Flow Cytometry Shared Core at the Jennie Smoly Caruthers Biotechnology Building (JSCBB) (RRID:SCR_019309) for technical support. This research was supported in part by the NIH/NCATS Colorado CTSA Grant Number UM1 TR004399. This research was supported by The Shurl and Kay Curci Foundation (JQ).

## Funding

This work was supported by the Colorado Nutrition Obesity Research Center (NORC) Pilot & Feasibility Program Award (P30DK048250), NIH grant R01DK126710, the National Dairy Council, and Mead Johnson Nutrition. Contents are the authors’ sole responsibility and do not necessarily represent official NIH views.

## Author information

**Jing Qian**

**Parsa Ghadermazi**

**Soren Maret**

**Jennifer Kemp**

**Daniel Frank**

**Edward Melanson**

**Audrey Hendricks**

**Nancy Krebs**

**Minghua Tang**

**Matthew R. Olm**

## Affiliation

**Jing Qian, Parsa Ghadermazi, Matthew R. Olm**

Department of Integrative Physiology, University of Colorado Boulder, Boulder, CO, USA

**Soren Maret**

Molecular, Cellular, and Developmental Biology Department, University of Colorado Boulder, Boulder, CO, USA

**Jennifer F. Kemp, Nancy Krebs**

Department of Pediatrics, Section of Nutrition, University of Colorado School of Medicine, Aurora, CO, USA

**Daniel Frank**

Department of Medicine-Infectious Disease, University of Colorado School of Medicine, Aurora, CO, USA

**Edward Melanson**

Division of Endocrinology, Metabolism, and Diabetes, University of Colorado Anschutz, Aurora, CO, USA

**Audrey Hendricks**

Department of Biomedical Informatics, University of Colorado Anschutz Medical Campus, Aurora, Colorado, USA

**Minghua Tang**

Department of Food Science and Human Nutrition, Colorado State University, Fort Collins, CO 80526, USA

## Contributions

M.R.O. conceived and supervised the study and secured funding. J.Q. and S.M. performed MIg-Seq experiments. J.Q. performed DNA extraction, metagenomic sequencing, data analysis, and statistical analysis, and wrote the manuscript. P.G. developed the bioinformatic analysis pipeline. J.K., D.F., E.M., A.H., N.K., and M.T. designed the clinical trial and collected samples. M.R.O. edited the manuscript. All authors reviewed and approved the final manuscript.

## Ethics declarations

### Ethics approval and consent to participate

Infant fecal samples were collected as part of a randomized controlled trial registered at ClinicalTrials.gov (NCT05012930). Ethical approval was obtained through the Colorado Multiple Institutional Review Board (COMIRB 20-2232). Written informed consent was provided by all caregivers prior to enrollment.

### Competing interests

The authors declare no competing interests.

### Supplementary information

See supplementary material.

## Notes

### Competing Interest Statement

The authors have declared no competing interest.

## Reference

1. Tamburini, S., Shen, N., Wu, H. C. & Clemente, J. C. The microbiome in early life: implications for health outcomes. Nature Medicine 22, 713–722 (2016).

2. Sonnenburg, E. D. & Sonnenburg, J. L. The ancestral and industrialized gut microbiota and implications for human health. Nature Reviews Microbiology 17, 383–390 (2019).

3. Blaser, M. J. The theory of disappearing microbiota and the epidemics of chronic diseases. Nat Rev Immunol 17, 461–463 (2017).

4. Donaldson, G. P. et al. Gut microbiota utilize immunoglobulin A for mucosal colonization. Science 360, 795–800 (2018).

5. IgA and the intestinal microbiota: the importance of being specific. Mucosal Immunology 13, 12–21 (2020).

6. Moor, K. et al. High-avidity IgA protects the intestine by enchaining growing bacteria. Nature 544, 498–502 (2017).

7. Rio-Aige, K. et al. The Breast Milk Immunoglobulinome. Nutrients 13, (2021).

8. Donald, K., Petersen, C., Turvey, S. E., Finlay, B. B. & Azad, M. B. Secretory IgA: Linking microbes, maternal health, and infant health through human milk. Cell Host Microbe 30, 650–659 (2022).

9. Rognum, T. O., Thrane, S., Stoltenberg, L., Vege, A. & Brandtzaeg, P. Development of intestinal mucosal immunity in fetal life and the first postnatal months. Pediatr Res 32, 145–149 (1992).

10. Rogosch, T. et al. IgA response in preterm neonates shows little evidence of antigen-driven selection. J Immunol 189, 5449–5456 (2012).

11. Planer, J. et al. Development of the gut microbiota and mucosal IgA responses in twins and gnotobiotic mice. Nature 534, 263–266 (2016).

12. Zheng, W. et al. Microbiota-targeted maternal antibodies protect neonates from enteric infection. Nature 577, 543–548 (2020).

13. Al Nabhani, Z. et al. A Weaning Reaction to Microbiota Is Required for Resistance to Immunopathologies in the Adult. Immunity 50, 1276–1288.e5 (2019).

14. Stewart, C. J. et al. Temporal development of the gut microbiome in early childhood from the TEDDY study. Nature 562, 583–588 (2018).

15. Palm, N. W. et al. Immunoglobulin A coating identifies colitogenic bacteria in inflammatory bowel disease. Cell 158, 1000–1010 (2014).

16. van Gogh, M. et al. Next-generation IgA-SEQ allows for high-throughput, anaerobic, and metagenomic assessment of IgA-coated bacteria. Microbiome 12, 211 (2024).

17. Olm, M. R., Spencer, S. P., Takeuchi, T., Silva, E. L. & Sonnenburg, J. L. Metagenomic immunoglobulin sequencing reveals IgA coating of microbial strains in the healthy human gut. Nat Microbiol 10, 112–125 (2025).

18. Viladomiu, M. et al. IgA-coated enriched in Crohn’s disease spondyloarthritis promote T17-dependent inflammation. Sci Transl Med 9, (2017).

19. Tang, M. et al. Effects of Complementary Feeding With Different Protein-Rich Foods on Infant Growth and Gut Health: Study Protocol. Front Pediatr 9, 793215 (2021).

20. Di Tommaso, P. et al. Nextflow enables reproducible computational workflows. Nature Biotechnology 35, 316–319 (2017).

21. Chen, S., Zhou, Y., Chen, Y. & Gu, J. fastp: an ultra-fast all-in-one FASTQ preprocessor. Bioinformatics 34, i884–i890 (2018).

22. Langmead, B. & Salzberg, S. L. Fast gapped-read alignment with Bowtie 2. Nature Methods 9, 357–359 (2012).

23. Shaw, J. & Yu, Y. W. Rapid species-level metagenome profiling and containment estimation with sylph. Nature Biotechnology 43, 1348–1359 (2024).

24. Parks, D. H. et al. GTDB: an ongoing census of bacterial and archaeal diversity through a phylogenetically consistent, rank normalized and complete genome-based taxonomy. Nucleic Acids Res 50, D785–D794 (2022).

25. Lozupone, C. & Knight, R. UniFrac: a New Phylogenetic Method for Comparing Microbial Communities. Applied and Environmental Microbiology (2005) doi:10.1128/AEM.71.12.8228-8235.2005.

26. McMurdie, P. J. & Holmes, S. phyloseq: An R Package for Reproducible Interactive Analysis and Graphics of Microbiome Census Data. PLOS ONE 8, e61217 (2013).

27. Anderson, M. J. A new method for non-parametric multivariate analysis of variance. Austral Ecology 26, 32–46 (2001).

28. Forestell, C. A. et al. Effects of early exposure to protein-rich complementary foods on toddlers liking and acceptance: The MINT study. Curr. Dev. Nutr. 9, 106808 (2025).

29. Bunker, J. J. et al. Natural polyreactive IgA antibodies coat the intestinal microbiota. Science 358, (2017).

30. Rollenske, T. et al. Parallelism of intestinal secretory IgA shapes functional microbial fitness. Nature 598, 657–661 (2021).

31. Sela, D. A. et al. The genome sequence of Bifidobacterium longum subsp. infantis reveals adaptations for milk utilization within the infant microbiome. Proc Natl Acad Sci U S A 105, 18964–18969 (2008).

32. Milani, C. et al. The First Microbial Colonizers of the Human Gut: Composition, Activities, and Health Implications of the Infant Gut Microbiota. Microbiol Mol Biol Rev 81, (2017).

33. Peterson, D. A., McNulty, N. P., Guruge, J. L. & Gordon, J. I. IgA response to symbiotic bacteria as a mediator of gut homeostasis. Cell Host Microbe 2, 328–339 (2007).

34. Crost, E. H. et al. The mucin-degradation strategy of Ruminococcus gnavus: The importance of intramolecular trans-sialidases. Gut Microbes 7, 302–312 (2016).

35. Henke, M. T. et al. , a member of the human gut microbiome associated with Crohn’s disease, produces an inflammatory polysaccharide. Proc Natl Acad Sci U S A 116, 12672–12677 (2019).

36. Bäckhed, F. et al. Dynamics and Stabilization of the Human Gut Microbiome during the First Year of Life. Cell Host Microbe 17, 852 (2015).

37. Bokulich, N. A. et al. Antibiotics, birth mode, and diet shape microbiome maturation during early life. Sci Transl Med 8, 343ra82 (2016).

38. Gensollen, T., Iyer, S. S., Kasper, D. L. & Blumberg, R. S. How colonization by microbiota in early life shapes the immune system. Science 352, 539–544 (2016).

39. Yatsunenko, T. et al. Human gut microbiome viewed across age and geography. Nature 486, 222–227 (2012).

40. Gopalakrishna, K. P. et al. Maternal IgA protects against the development of necrotizing enterocolitis in preterm infants. Nat Med 25, 1110–1115 (2019).

41. Macpherson, A. J., de Agüero, M. G. & Ganal-Vonarburg, S. C. How nutrition and the maternal microbiota shape the neonatal immune system. Nat Rev Immunol 17, 508–517 (2017).

42. Koch, M. A. et al. Maternal IgG and IgA Antibodies Dampen Mucosal T Helper Cell Responses in Early Life. Cell 165, 827–841 (2016).

43. Donald, K. et al. Human milk IgA promotes normal immune development by limiting Th17-inducing in the infant gut. Proc Natl Acad Sci U S A 122, e2501030122 (2025).

44. Rogier, E. W. et al. Secretory antibodies in breast milk promote long-term intestinal homeostasis by regulating the gut microbiota and host gene expression. Proc Natl Acad Sci U S A 111, 3074–3079 (2014).

45. Torow, N., Marsland, B. J., Hornef, M. W. & Gollwitzer, E. S. Neonatal mucosal immunology. Mucosal Immunol 10, 5–17 (2017).

46. Henrick, B. M. et al. Bifidobacteria-mediated immune system imprinting early in life. Cell 184, 3884–3898.e11 (2021).

47. Kim, M., Qie, Y., Park, J. & Kim, C. H. Gut Microbial Metabolites Fuel Host Antibody Responses. Cell Host Microbe 20, 202–214 (2016).

48. Takeuchi, T. et al. Acetate differentially regulates IgA reactivity to commensal bacteria. Nature 595, 560–564 (2021).

49. Johansen, F.-E. & Kaetzel, C. S. Regulation of the polymeric immunoglobulin receptor and IgA transport: new advances in environmental factors that stimulate pIgR expression and its role in mucosal immunity. Mucosal Immunol 4, 598–602 (2011).

50. Makki, K., Deehan, E. C., Walter, J. & Bäckhed, F. The Impact of Dietary Fiber on Gut Microbiota in Host Health and Disease. Cell Host Microbe 23, 705–715 (2018).

51. Olm, M. R. et al. Robust variation in infant gut microbiome assembly across a spectrum of lifestyles. Science 376, 1220–1223 (2022).

