## Supplementary material for "IgA Targeting in the Infant Gut Is Modulated by Diet and Increasingly Directed Towards Persistent Species"

**Supplementary Table S1.** PERMANOVA results for β-diversity comparisons across age groups and timepoints

| **Distance** | **Comparison** | **n** | **R2** | **F_stat** | **P** |
| --- | --- | --- | --- | --- | --- |
| **Unweighted — IgA+** | 6mo vs 12mo (infant only, within-subject permutation) | 32 | 0.0564 | 1.79 | 0.007 |
| **Unweighted — Native** | 6mo vs 12mo (infant only, within-subject permutation) | 32 | 0.0465 | 1.46 | 0.074 |
| **Weighted — IgA+** | 6mo vs 12mo (infant only, within-subject permutation) | 32 | 0.0617 | 1.97 | 0.028 |
| **Weighted — Native** | 6mo vs 12mo (infant only, within-subject permutation) | 32 | 0.0437 | 1.37 | 0.148 |
| **Unweighted — IgA+** | Infant vs Adult | 62 | 0.285 | 23.9 | 0.001 |
| **Unweighted — Native** | Infant vs Adult | 62 | 0.329 | 29.4 | 0.001 |
| **Weighted — IgA+** | Infant vs Adult | 62 | 0.326 | 29.1 | 0.001 |
| **Weighted — Native** | Infant vs Adult | 62 | 0.345 | 31.7 | 0.001 |

**Supplementary Table S2.** Wilcoxon rank-sum test results for within- and between-subject UniFrac distances in adults and infants (unweighted UniFrac; native and IgA-enriched fractions)

| **Distance** | **Comparison** | **n_a** | **median_a** | **n_b** | **median_b** | **P** |
| --- | --- | --- | --- | --- | --- | --- |
| **Unweighted — Native** | Adult: within vs between subject | 11 | 0.294 | 424 | 0.569 | 1.16E-07 |
| **Unweighted — Native** | Infant: within vs between subject | 16 | 0.521 | 480 | 0.536 | 0.336 |
| **Unweighted — Native** | Within-subject: Adult vs Infant temporal distance | 11 | 0.294 | 16 | 0.521 | 0.00392 |
| **Unweighted — Native** | Infant: within vs between (6mo-6mo) | 16 | 0.521 | 120 | 0.513 | 0.879 |
| **Unweighted — Native** | Infant: within vs between (12mo-12mo) | 16 | 0.521 | 120 | 0.554 | 0.176 |
| **Unweighted — Native** | Infant: within vs between (6mo-12mo) | 16 | 0.521 | 240 | 0.542 | 0.267 |
| **Unweighted — IgA+** | Adult: within vs between subject | 11 | 0.333 | 424 | 0.582 | 2.26E-08 |
| **Unweighted — IgA+** | Infant: within vs between subject | 16 | 0.505 | 480 | 0.548 | 0.0403 |
| **Unweighted — IgA+** | Within-subject: Adult vs Infant temporal distance | 11 | 0.333 | 16 | 0.505 | 7.78E-04 |
| **Unweighted — IgA+** | Infant: within vs between (6mo-6mo) | 16 | 0.505 | 120 | 0.521 | 0.269 |
| **Unweighted — IgA+** | Infant: within vs between (12mo-12mo) | 16 | 0.505 | 120 | 0.564 | 0.0294 |
| **Unweighted — IgA+** | Infant: within vs between (6mo-12mo) | 16 | 0.505 | 240 | 0.561 | 0.0196 |

**Supplementary Table S3.** Linear mixed-effects model results for IgA fold-change comparisons of transient versus persistent taxa and developmental changes in Bifidobacterium IgA targeting

| **Panel** | **Comparison** | **n_obs** | **n_subjects** | **median_FC** | **mean_FC** | **LMM_Estimate** | **p_value** |
| --- | --- | --- | --- | --- | --- | --- | --- |
| a | Transient vs Persistent (6mo) | 454 | 16 | 0.88 vs 0.2 | 0.87 vs -0.12 | 0.4278 | 0.4728 |
| b | Persistent taxa: 6mo vs 12mo | 398 | 16 | 0.2 → 0.59 | -0.12 → 2 | 2.1235 | 0.0000 |
| c | Persistent Bifido: 6mo vs 12mo | 84 | 14 | -0.09 → 2.29 | 0.06 → 3.96 | 3.8951 | 0.0001 |
| d | Bifido — Meat: 6mo vs 12mo | 40 | 6 | -0.19 → 2.4 | 0.03 → 4.13 | 4.1033 | 0.0006 |
| e | Bifido — Plant: 6mo vs 12mo | 34 | 6 | -0.17 → 2.65 | 0.04 → 4.74 | 4.7001 | 0.0249 |
| f | Bifido — Human milk: 6mo vs 12mo | 28 | 6 | -0.9 → 3.26 | -0.97 → 4.49 | 5.4579 | 0.0111 |
| g | Bifido — Formula: 6mo vs 12mo | 56 | 8 | 0.39 → 1.81 | 0.58 → 3.69 | 3.1136 | 0.0042 |

**Supplementary Table S4** Phylum-level IgA fold-change by clinical covariates at enrollment (Wilcoxon rank-sum test, BH-corrected)

| **Covariate** | **Phylum** | **Group 1** | **Group 2** | **Median (Group 1)** | **Median (Group 2)** | **p-value** | **q-value** |
| --- | --- | --- | --- | --- | --- | --- | --- |
| **Childcare attendance** | Actinomycetota | No (n=11) | Yes (n=4) | 0.284 | -0.447 | 0.1770 | 0.5000 |
| **Childcare attendance** | Bacillota | No (n=10) | Yes (n=4) | -0.192 | 1.500 | 0.2400 | 0.5000 |
| **Childcare attendance** | Pseudomonadota | No (n=11) | Yes (n=3) | 0.517 | 3.770 | 0.2740 | 0.5000 |
| **Childcare attendance** | Bacillota C | No (n=5) | Yes (n=4) | 3.600 | 1.310 | 0.2860 | 0.5000 |
| **Childcare attendance** | Bacteroidota | No (n=9) | Yes (n=4) | 0.082 | -0.603 | 0.7100 | 0.9930 |
| **Childcare attendance** | Bacillota A | No (n=11) | Yes (n=4) | 0.546 | 0.717 | 0.8510 | 0.9930 |
| **Childcare attendance** | Bacillota I | No (n=11) | Yes (n=4) | 0.022 | 1.340 | 1.000 | 1.000 |
| **Delivery mode** | Bacillota C | Vaginal (n=6) | CSection (n=3) | 1.310 | 3.600 | 0.2620 | 0.9640 |
| **Delivery mode** | Bacillota | Vaginal (n=8) | CSection (n=6) | 1.360 | -0.192 | 0.4910 | 0.9640 |
| **Delivery mode** | Bacteroidota | Vaginal (n=8) | CSection (n=5) | 0.077 | -0.380 | 0.5240 | 0.9640 |
| **Delivery mode** | Bacillota I | Vaginal (n=9) | CSection (n=6) | -0.060 | 0.563 | 0.6070 | 0.9640 |
| **Delivery mode** | Bacillota A | Vaginal (n=9) | CSection (n=6) | 0.944 | 0.525 | 0.6890 | 0.9640 |
| **Delivery mode** | Pseudomonadota | Vaginal (n=8) | CSection (n=6) | 1.390 | 1.580 | 0.8460 | 0.9870 |
| **Delivery mode** | Actinomycetota | Vaginal (n=9) | CSection (n=6) | 0.213 | -0.236 | 1.000 | 1.000 |
| **Sex** | Actinomycetota | Female (n=10) | Male (n=5) | -0.418 | 0.718 | 0.1650 | 0.6590 |
| **Sex** | Pseudomonadota | Female (n=9) | Male (n=5) | 2.640 | 0.000 | 0.3490 | 0.6590 |
| **Sex** | Bacillota A | Female (n=10) | Male (n=5) | 0.235 | 0.944 | 0.3710 | 0.6590 |
| **Sex** | Bacillota I | Female (n=10) | Male (n=5) | 0.563 | -0.130 | 0.4400 | 0.6590 |
| **Sex** | Bacillota | Female (n=10) | Male (n=4) | 0.129 | 1.140 | 0.8390 | 0.9370 |
| **Sex** | Bacteroidota | Female (n=10) | Male (n=3) | -0.154 | 0.082 | 0.9370 | 0.9370 |

**Supplementary Table S5** Phylum-level IgA fold-change across dietary arms at enrollment (Kruskal-Wallis test, BH-corrected)

| **Phylum** | **Median (Meat)** | **Median (Dairy)** | **Median (Plant)** | **Median (Reference)** | **p-value** | **q-value** |
| --- | --- | --- | --- | --- | --- | --- |
| **Bacillota** | 2.29 | 2.56 | 0.219 | -1.87 | 0.4290 | 0.7530 |
| **Actinomycetota** | 0.248 | -1.25 | 0.318 | -0.644 | 0.4950 | 0.7530 |
| **Bacillota I** | 0.144 | 3.11 | 0.4 | -1.7 | 0.5500 | 0.7530 |
| **Bacteroidota** | -0.38 | 1.62 | 0.0725 | NA | 0.6660 | 0.7530 |
| **Pseudomonadota** | 3.26 | 4.76 | 0.974 | 0.517 | 0.6660 | 0.7530 |
| **Verrucomicrobiota** | 0.452 | 4.14 | 0.0177 | NA | 0.6670 | 0.7530 |
| **Bacillota C** | 2.38 | 3.68 | 2.33 | NA | 0.7410 | 0.7530 |
| **Bacillota A** | 0.235 | 2.39 | 1.01 | 0.504 | 0.7530 | 0.7530 |

**Supplemental S6** Species-level IgA fold-change by feeding mode at enrollment (Wilcoxon rank-sum test, BH-corrected)

| **Species** | **n (Human milk)** | **n (Formula)** | **Median FC (Human milk)** | **Median FC (Formula)** | **p-value** | **q-value** |
| --- | --- | --- | --- | --- | --- | --- |
| **B. longum** | 6 | 8 | -1.410 | -0.374 | 0.04260 | 0.2490 |
| **Parabacteroides distasonis** | 3 | 4 | 6.510 | -4.970 | 0.04970 | 0.2490 |
| **P. vulgatus** | 3 | 4 | -10.000 | 1.100 | 0.1080 | 0.3610 |
| **Clostridium innocuum** | 3 | 4 | -0.060 | 4.360 | 0.2290 | 0.4820 |
| **T. ramosa** | 5 | 8 | -0.130 | 0.815 | 0.2410 | 0.4820 |
| **Bifidobacterium breve** | 4 | 6 | 3.080 | 0.938 | 0.3520 | 0.5870 |
| **F. plautii** | 4 | 7 | 0.825 | 0.347 | 0.4120 | 0.5890 |
| **E. lenta** | 6 | 6 | -0.822 | -0.020 | 0.4850 | 0.6060 |
| **Escherichia coli** | 6 | 8 | 0.980 | 2.240 | 0.6050 | 0.6720 |
| **R. gnavus** | 5 | 9 | -0.509 | 0.251 | 0.8980 | 0.8980 |

**Supplemental S7** Phylum- and species-level IgA fold-change across dietary arms at 12 months (Kruskal-Wallis test, BH-corrected)

**Section 1** Table_12mo_arm_phylum_KW

| **Phylum** | **Median FC (Meat)** | **Median FC (Dairy)** | **Median FC (Plant)** | **Median FC (Reference)** | **p-value** | **q-value** |
| --- | --- | --- | --- | --- | --- | --- |
| **Verrucomicrobiota** | 2.01 | NA | 0.409 | NA | 0.2480 | 0.8170 |
| **Bacillota C** | 1.12 | 2.29 | 0.594 | -0.665 | 0.3460 | 0.8170 |
| **Pseudomonadota** | 1.8 | -0.528 | 0.981 | 1.09 | 0.5400 | 0.8170 |
| **Bacillota I** | 0.373 | 0.199 | -1.15 | -0.523 | 0.5490 | 0.8170 |
| **Bacillota A** | 0.93 | 0.0613 | 1.09 | 0.183 | 0.5820 | 0.8170 |
| **Bacillota** | 0.758 | 0.00218 | 1.76 | -1.13 | 0.6950 | 0.8170 |
| **Actinomycetota** | 0.628 | 0.209 | 0.482 | -0.593 | 0.7750 | 0.8170 |
| **Bacteroidota** | 0.361 | -0.872 | 0.0999 | NA | 0.8170 | 0.8170 |

**Section 2** Table_12mo_arm_species_KW

| **Species** | **Median FC (Meat)** | **Median FC (Dairy)** | **Median FC (Plant)** | **Median FC (Reference)** | **p-value** | **q-value** |
| --- | --- | --- | --- | --- | --- | --- |
| **B. longum** | 0.984 | -0.376 | -0.595 | NA | 0.1960 | 0.9640 |
| **Bifidobacterium breve** | 0.548 | -0.333 | 6.32 | NA | 0.5060 | 0.9640 |
| **E. lenta** | -0.394 | -0.517 | -2.15 | -1.29 | 0.5130 | 0.9640 |
| **Escherichia coli** | 1.44 | -0.528 | 1.14 | 1.09 | 0.6700 | 0.9640 |
| **F. plautii** | 0.506 | -0.625 | 0.475 | -1.1 | 0.7170 | 0.9640 |
| **T. ramosa** | -0.227 | 0.469 | -1.44 | -0.281 | 0.7320 | 0.9640 |
| **Anaerostipes hadrus** | 1.75 | 2.08 | 5.91 | NA | 0.7550 | 0.9640 |
| **B. bifidum** | 2.24 | 2.5 | 0.764 | NA | 0.7570 | 0.9640 |
| **P. vulgatus** | 0.626 | -1.32 | -0.434 | NA | 0.8110 | 0.9640 |
| **Clostridium innocuum** | 0.871 | -0.0712 | 1.11 | -0.766 | 0.9430 | 0.9640 |
| **R. gnavus** | 0.0363 | 0.014 | 0.489 | 0.594 | 0.9640 | 0.9640 |

**Supplementary Table S8** Nutrient–species IgA fold-change associations meeting q < 0.10 at 12 months (Spearman's ρ, BH-corrected within each nutrient)

| **Species** | **Nutrient** | **N** | **ρ** | **p-value** | **q-value** |
| --- | --- | --- | --- | --- | --- |
| **B. longum** | Cholesterol | 11 | 0.909 | <0.001 | 0.001000 |
| **F. plautii** | Alpha-carotene | 12 | 0.825 | 0.001000 | 0.01000 |
| **T. ramosa** | Trans fat (18:1) | 14 | 0.758 | 0.001700 | 0.01800 |
| **R. gnavus** | GLA (18:3 n-6) | 13 | 0.729 | 0.004700 | 0.05200 |
| **E. lenta** | Erythritol | 12 | 0.746 | 0.005400 | 0.05900 |
| **E. lenta** | CLA (18:2, total) | 12 | 0.662 | 0.01900 | 0.07000 |
| **B. bifidum** | Synthetic folic acid | 9 | -0.810 | 0.008100 | 0.08900 |
| **T. ramosa** | Butyric acid (SFA 4:0) | 14 | 0.589 | 0.02650 | 0.09600 |

**Supplementary Table S9** Nutrient–species native relative abundance associations meeting q < 0.10 at 12 months (Spearman's ρ, BH-corrected within each nutrient; 1,683 total tests)

| **Nutrient** | **Species** | **N** | **ρ** | **p-value** | **q-value** |
| --- | --- | --- | --- | --- | --- |
| **Myristic acid (SFA 14:0)** | B. longum | 14 | -0.717 | 0.003870 | 0.04260 |
| **Lauric acid (SFA 12:0)** | B. longum | 14 | -0.713 | 0.004200 | 0.04620 |
| **Erythritol** | E. lenta | 14 | -0.711 | 0.004330 | 0.04770 |
| **Solid Fats** | B. longum | 14 | -0.703 | 0.005050 | 0.05550 |
| **Caprylic acid (SFA 8:0)** | B. longum | 14 | -0.656 | 0.01080 | 0.06130 |
| **Caprylic acid (SFA 8:0)** | F. plautii | 14 | 0.654 | 0.01120 | 0.06130 |
| **Erythritol** | T. ramosa | 14 | -0.654 | 0.01120 | 0.06130 |
| **Genistein** | F. plautii | 14 | 0.676 | 0.007920 | 0.08710 |
| **Erythritol** | Clostridium innocuum | 14 | -0.590 | 0.02630 | 0.09660 |

**Supplementary Table S10** Nutrient–species IgA fold-change and native relative abundance associations at 12 months: Spearman's ρ for all pairs significant in at least one analysis (q < 0.10), with direction of effect compared across analyses (BH-corrected within each nutrient; 1,683 tests per analysis)

| **Category** | **Nutrient** | **Species** | **ρ (FC)** | **q (FC)** | **ρ (RA)** | **q (RA)** | **Direction** |
| --- | --- | --- | --- | --- | --- | --- | --- |
| **FC and RA significant** | Solid Fats | B. longum | 0.780 | 0.05110 | -0.703 | 0.05550 | Opposite |
| **FC and RA significant** | Erythritol | E. lenta | 0.746 | 0.05900 | -0.711 | 0.04770 | Opposite |
| **FC and RA significant** | Erythritol | T. ramosa | 0.628 | 0.08840 | -0.654 | 0.06130 | Opposite |
| **FC and RA significant** | Erythritol | Clostridium innocuum | 0.638 | 0.08920 | -0.590 | 0.09660 | Opposite |
| **FC significant only** | Cholesterol | B. longum | 0.909 | 0.001160 | -0.528 | 0.3000 | Opposite |
| **FC significant only** | CLA (trans-10, cis-12) | B. longum | 0.874 | 0.004760 | -0.561 | 0.4060 | Opposite |
| **FC significant only** | Alpha-carotene | F. plautii | 0.825 | 0.01050 | 0.042 | 0.9180 | Same |
| **FC significant only** | Animal protein | B. longum | 0.836 | 0.01470 | -0.333 | 0.6710 | Opposite |
| **FC significant only** | EPA (20:5 n-3) | B. longum | 0.828 | 0.01810 | -0.413 | 0.3900 | Opposite |
| **FC significant only** | Trans fat (18:1) | T. ramosa | 0.758 | 0.01840 | -0.437 | 0.6480 | Opposite |
| **FC significant only** | Beta-carotene | F. plautii | 0.720 | 0.04530 | 0.205 | 0.8990 | Same |
| **FC significant only** | Beta-carotene | R. gnavus | 0.703 | 0.04530 | -0.293 | 0.8520 | Opposite |
| **FC significant only** | CLA (18:2, total) | B. longum | 0.770 | 0.04570 | -0.506 | 0.7130 | Opposite |
| **FC significant only** | CLA (18:2, total) | T. ramosa | 0.673 | 0.04570 | -0.376 | 0.7840 | Opposite |
| **FC significant only** | GLA (18:3 n-6) | R. gnavus | 0.729 | 0.05160 | -0.182 | 0.9330 | Opposite |
| **FC significant only** | CLA cis 9. trans 11 | B. longum | 0.733 | 0.05610 | -0.502 | 0.7440 | Opposite |
| **FC significant only** | CLA cis 9. trans 11 | T. ramosa | 0.667 | 0.05610 | -0.356 | 0.7530 | Opposite |
| **FC significant only** | Trans fat (18:1) | B. longum | 0.727 | 0.06160 | -0.647 | 0.1370 | Opposite |
| **FC significant only** | Trans fat (18:1) | E. lenta | 0.671 | 0.06170 | -0.163 | 0.8980 | Opposite |
| **FC significant only** | CLA (18:2, total) | E. lenta | 0.662 | 0.06970 | -0.174 | 0.7840 | Opposite |
| **FC significant only** | Total Trans Fatty Acids TRANS | T. ramosa | 0.688 | 0.07190 | -0.420 | 0.7430 | Opposite |
| **FC significant only** | PUFA 20.4 arachidonic acid. undifferentiated | B. longum | 0.761 | 0.07200 | -0.539 | 0.5130 | Opposite |
| **FC significant only** | Vitamin A (total, IU) | R. gnavus | 0.709 | 0.07350 | -0.334 | 0.7870 | Opposite |
| **FC significant only** | CLA cis 9. trans 11 | E. lenta | 0.657 | 0.07400 | -0.188 | 0.7530 | Opposite |
| **FC significant only** | Biochanin A (isoflavone) | B. longum | -0.758 | 0.07540 | 0.435 | 0.4200 | Opposite |
| **FC significant only** | Formononetin (isoflavone) | B. longum | -0.758 | 0.07540 | 0.435 | 0.4200 | Opposite |
| **FC significant only** | Beta-carotene equivalents | F. plautii | 0.685 | 0.07650 | 0.227 | 0.8280 | Same |
| **FC significant only** | Beta-carotene equivalents | R. gnavus | 0.703 | 0.07650 | -0.277 | 0.8280 | Opposite |
| **FC significant only** | Total Trans Fatty Acids TRANS | B. longum | 0.691 | 0.08080 | -0.603 | 0.2480 | Opposite |
| **FC significant only** | Total Trans Fatty Acids TRANS | E. lenta | 0.650 | 0.08080 | -0.154 | 0.9000 | Opposite |
| **FC significant only** | Caproic acid (SFA 6:0) | P. vulgatus | 0.815 | 0.08170 | -0.474 | 0.4780 | Opposite |
| **FC significant only** | PUFA 22.5 n 3 docosapentaenoic acid .DPA. | B. bifidum | -0.770 | 0.08350 | 0.533 | 0.1820 | Opposite |
| **FC significant only** | PUFA 22.5 n 3 docosapentaenoic acid .DPA. | B. longum | 0.734 | 0.08350 | -0.349 | 0.6080 | Opposite |
| **FC significant only** | Alpha-carotene | R. gnavus | 0.654 | 0.08440 | -0.378 | 0.6170 | Opposite |
| **FC significant only** | Trans fat (18:1) | B. bifidum | -0.712 | 0.08640 | 0.336 | 0.8810 | Opposite |
| **FC significant only** | Total Trans Fatty Acids TRANS | B. bifidum | -0.712 | 0.08640 | 0.217 | 0.9000 | Opposite |
| **FC significant only** | Synthetic folic acid | B. bifidum | -0.810 | 0.08890 | 0.039 | 0.9840 | Opposite |
| **FC significant only** | Erythritol | B. bifidum | -0.709 | 0.08920 | 0.224 | 0.6060 | Opposite |
| **FC significant only** | Synthetic Alpha Tocopherol all rac alpha tocopherol or dl alpha tocopherol | B. bifidum | -0.687 | 0.08990 | 0.234 | 0.7480 | Opposite |
| **FC significant only** | Synthetic Alpha Tocopherol all rac alpha tocopherol or dl alpha tocopherol | Clostridium innocuum | 0.616 | 0.08990 | -0.384 | 0.5600 | Opposite |
| **FC significant only** | Synthetic Alpha Tocopherol all rac alpha tocopherol or dl alpha tocopherol | E. lenta | 0.664 | 0.08990 | -0.428 | 0.5600 | Opposite |
| **FC significant only** | Synthetic Alpha Tocopherol all rac alpha tocopherol or dl alpha tocopherol | F. plautii | 0.620 | 0.08990 | -0.191 | 0.7480 | Opposite |
| **FC significant only** | Synthetic Alpha Tocopherol all rac alpha tocopherol or dl alpha tocopherol | T. ramosa | 0.580 | 0.08990 | -0.415 | 0.5600 | Opposite |
| **FC significant only** | Butyric acid (SFA 4:0) | E. lenta | 0.690 | 0.09410 | -0.106 | 0.7900 | Opposite |
| **FC significant only** | Butyric acid (SFA 4:0) | P. vulgatus | 0.762 | 0.09410 | -0.318 | 0.7900 | Opposite |
| **FC significant only** | Butyric acid (SFA 4:0) | B. longum | 0.638 | 0.09550 | -0.441 | 0.7900 | Opposite |
| **FC significant only** | Butyric acid (SFA 4:0) | T. ramosa | 0.589 | 0.09550 | -0.157 | 0.7900 | Opposite |
| **FC significant only** | Alpha-carotene | B. bifidum | -0.712 | 0.09850 | 0.083 | 0.9180 | Opposite |
| **FC significant only** | Alpha-carotene | Clostridium innocuum | 0.608 | 0.09850 | -0.360 | 0.6170 | Opposite |
| **FC significant only** | GLA (18:3 n-6) | B. bifidum | -0.758 | 0.09940 | 0.034 | 0.9340 | Opposite |
| **RA significant only** | Myristic acid (SFA 14:0) | B. longum | 0.555 | 0.2110 | -0.717 | 0.04260 | Opposite |
| **RA significant only** | Lauric acid (SFA 12:0) | B. longum | 0.309 | 0.5580 | -0.713 | 0.04620 | Opposite |
| **RA significant only** | Caprylic acid (SFA 8:0) | B. longum | 0.300 | 0.6890 | -0.656 | 0.06130 | Opposite |
| **RA significant only** | Caprylic acid (SFA 8:0) | F. plautii | -0.151 | 0.7040 | 0.654 | 0.06130 | Opposite |
| **RA significant only** | Genistein | F. plautii | -0.063 | 0.9690 | 0.676 | 0.08710 | Opposite |

**Supplementary Table S11** Spearman correlations between 153 dietary nutrients and genus-level Bifidobacterium IgA fold-change (n = 14)

| **Nutrient** | **N** | **ρ** | **p-value** | **q-value** |
| --- | --- | --- | --- | --- |
| **Alpha-carotene** | 14 | -0.684 | 0.007040 | 0.5270 |
| **Vitamin A (total, IU)** | 14 | -0.648 | 0.01210 | 0.5270 |
| **Beta-carotene equivalents** | 14 | -0.644 | 0.01290 | 0.5270 |
| **Beta-carotene** | 14 | -0.640 | 0.01380 | 0.5270 |
| **Vitamin A (RAE)** | 14 | -0.596 | 0.02460 | 0.7360 |
| **Vitamin A (RE)** | 14 | -0.582 | 0.02890 | 0.7360 |
| **Pectins** | 14 | -0.556 | 0.03890 | 0.8510 |
| **Erythritol** | 14 | -0.431 | 0.1240 | 1.000 |
| **Beta-cryptoxanthin** | 14 | -0.418 | 0.1370 | 1.000 |
| **Lauric acid (SFA 12:0)** | 14 | -0.407 | 0.1490 | 1.000 |
| **Vitamin D** | 14 | -0.402 | 0.1540 | 1.000 |
| **Vitamin D3 cholecalciferol** | 14 | -0.402 | 0.1540 | 1.000 |
| **Whole Grains ounce equivalents** | 14 | -0.392 | 0.1660 | 1.000 |
| **Mannitol** | 14 | -0.377 | 0.1830 | 1.000 |
| **Synthetic Alpha Tocopherol all rac alpha tocopherol or dl alpha tocopherol** | 14 | -0.364 | 0.2010 | 1.000 |
| **Synthetic folic acid** | 14 | -0.355 | 0.2120 | 1.000 |
| **Maltose** | 14 | -0.354 | 0.2150 | 1.000 |
| **Lariciresinol** | 14 | -0.354 | 0.2150 | 1.000 |
| **Lutein . Zeaxanthin** | 14 | -0.349 | 0.2210 | 1.000 |
| **Daidzein** | 14 | -0.343 | 0.2300 | 1.000 |
| **Myristic acid (SFA 14:0)** | 14 | -0.341 | 0.2330 | 1.000 |
| **Sorbitol** | 14 | -0.311 | 0.2790 | 1.000 |
| **Secoisolariciresinol** | 14 | -0.305 | 0.2880 | 1.000 |
| **Total Grains ounce equivalents** | 14 | -0.301 | 0.2960 | 1.000 |
| **Glucose** | 14 | -0.297 | 0.3030 | 1.000 |
| **Dietary fiber (soluble)** | 14 | -0.297 | 0.3030 | 1.000 |
| **Pantothenic Acid** | 14 | -0.292 | 0.3110 | 1.000 |
| **Trans fat (18:1)** | 14 | -0.288 | 0.3180 | 1.000 |
| **Natural Alpha Tocopherol RRR alpha tocopherol or d alpha tocopherol** | 14 | -0.288 | 0.3180 | 1.000 |
| **Matairesinol** | 14 | -0.288 | 0.3180 | 1.000 |
| **Total Trans Fatty Acids TRANS** | 14 | -0.279 | 0.3340 | 1.000 |
| **Pinoresinol** | 14 | -0.279 | 0.3340 | 1.000 |
| **Total Alpha Tocopherol Equivalents** | 14 | -0.275 | 0.3420 | 1.000 |
| **Vitamin E International Units** | 14 | -0.275 | 0.3420 | 1.000 |
| **Fructose** | 14 | -0.270 | 0.3500 | 1.000 |
| **MUFA 16.1 palmitoleic acid** | 14 | -0.270 | 0.3500 | 1.000 |
| **Total Lignans** | 14 | -0.270 | 0.3500 | 1.000 |
| **Vitamin E Total Alpha Tocopherol** | 14 | -0.266 | 0.3580 | 1.000 |
| **Iron** | 14 | -0.266 | 0.3580 | 1.000 |
| **Galactose** | 14 | 0.264 | 0.3620 | 1.000 |
| **GLA (18:3 n-6)** | 14 | -0.262 | 0.3650 | 1.000 |
| **Vitamin K phylloquinone** | 14 | -0.253 | 0.3830 | 1.000 |
| **Potassium** | 14 | -0.253 | 0.3830 | 1.000 |
| **Total sugars** | 14 | -0.248 | 0.3920 | 1.000 |
| **Lactose** | 14 | -0.246 | 0.3960 | 1.000 |
| **Genistein** | 14 | -0.241 | 0.4070 | 1.000 |
| **Vitamin C ascorbic acid** | 14 | -0.240 | 0.4090 | 1.000 |
| **PUFA 22.5 n 3 docosapentaenoic acid .DPA.** | 14 | -0.231 | 0.4270 | 1.000 |
| **Omega 3 Fatty Acids** | 14 | -0.226 | 0.4360 | 1.000 |
| **Vitamin D2 ergocalciferol** | 14 | -0.226 | 0.4380 | 1.000 |
| **Dietary fiber (insoluble)** | 14 | -0.218 | 0.4550 | 1.000 |
| **Capric acid (SFA 10:0)** | 14 | -0.213 | 0.4640 | 1.000 |
| **Manganese** | 14 | -0.209 | 0.4740 | 1.000 |
| **Caprylic acid (SFA 8:0)** | 14 | -0.205 | 0.4830 | 1.000 |
| **Total Carbohydrate** | 14 | -0.204 | 0.4830 | 1.000 |
| **Magnesium** | 14 | -0.204 | 0.4830 | 1.000 |
| **Starch** | 14 | -0.200 | 0.4930 | 1.000 |
| **Vitamin B 6 pyridoxine. pyridoxyl. . pyridoxamine** | 14 | -0.200 | 0.4930 | 1.000 |
| **Sucrose** | 14 | -0.196 | 0.5030 | 1.000 |
| **Betaine** | 14 | -0.191 | 0.5130 | 1.000 |
| **Lycopene** | 14 | -0.187 | 0.5230 | 1.000 |
| **Xylitol** | 14 | -0.186 | 0.5250 | 1.000 |
| **DHA (22:6 n-3)** | 14 | -0.183 | 0.5310 | 1.000 |
| **Copper** | 14 | -0.182 | 0.5330 | 1.000 |
| **Oxalic Acid** | 14 | -0.182 | 0.5330 | 1.000 |
| **Phytic Acid** | 14 | -0.178 | 0.5430 | 1.000 |
| **Dietary fiber (total)** | 14 | -0.174 | 0.5530 | 1.000 |
| **EPA (20:5 n-3)** | 14 | -0.171 | 0.5580 | 1.000 |
| **Caproic acid (SFA 6:0)** | 14 | -0.170 | 0.5610 | 1.000 |
| **Available Carbohydrate** | 14 | -0.169 | 0.5630 | 1.000 |
| **Niacin vitamin B3** | 14 | -0.152 | 0.6050 | 1.000 |
| **Calcium** | 14 | -0.152 | 0.6050 | 1.000 |
| **Biochanin A (isoflavone)** | 14 | 0.141 | 0.6310 | 1.000 |
| **Formononetin (isoflavone)** | 14 | 0.141 | 0.6310 | 1.000 |
| **Glycine** | 14 | 0.138 | 0.6370 | 1.000 |
| **X3 Methylhistidine** | 14 | -0.133 | 0.6510 | 1.000 |
| **Phenylalanine** | 14 | 0.130 | 0.6590 | 1.000 |
| **Histidine** | 14 | 0.130 | 0.6590 | 1.000 |
| **Gram Amount of Food** | 14 | -0.125 | 0.6700 | 1.000 |
| **Energy** | 14 | -0.125 | 0.6700 | 1.000 |
| **Energy kj** | 14 | -0.125 | 0.6700 | 1.000 |
| **Pinitol** | 14 | -0.124 | 0.6740 | 1.000 |
| **Solid Fats** | 14 | -0.121 | 0.6790 | 1.000 |
| **Vegetable protein** | 14 | -0.121 | 0.6810 | 1.000 |
| **SFA 20.0 arachidic acid** | 14 | 0.116 | 0.6920 | 1.000 |
| **Gamma Tocopherol** | 14 | -0.116 | 0.6920 | 1.000 |
| **PUFA 18.3 linolenic acid. undifferentiated** | 14 | -0.116 | 0.6920 | 1.000 |
| **Inositol** | 14 | -0.116 | 0.6920 | 1.000 |
| **Glycitein** | 14 | -0.115 | 0.6950 | 1.000 |
| **Added Sugars by Total sugars** | 14 | -0.114 | 0.6970 | 1.000 |
| **Coumestrol** | 14 | -0.110 | 0.7090 | 1.000 |
| **CLA (18:2, total)** | 14 | -0.108 | 0.7140 | 1.000 |
| **Beta Tocopherol** | 14 | -0.108 | 0.7140 | 1.000 |
| **Vitamin B12** | 14 | -0.108 | 0.7140 | 1.000 |
| **Selenium** | 14 | 0.108 | 0.7140 | 1.000 |
| **CLA cis 9. trans 11** | 14 | -0.106 | 0.7190 | 1.000 |
| **Butyric acid (SFA 4:0)** | 14 | -0.104 | 0.7240 | 1.000 |
| **Gluten** | 14 | -0.104 | 0.7240 | 1.000 |
| **Thiamin vitamin B1** | 14 | -0.103 | 0.7250 | 1.000 |
| **Zinc** | 14 | -0.103 | 0.7250 | 1.000 |
| **ALA (18:3 n-3)** | 14 | -0.103 | 0.7250 | 1.000 |
| **Proline** | 14 | 0.103 | 0.7250 | 1.000 |
| **Riboflavin vitamin B2** | 14 | -0.099 | 0.7370 | 1.000 |
| **Niacin Equivalents** | 14 | -0.095 | 0.7480 | 1.000 |
| **Valine** | 14 | 0.095 | 0.7480 | 1.000 |
| **Refined Grains ounce equivalents** | 14 | -0.092 | 0.7530 | 1.000 |
| **Delta Tocopherol** | 14 | 0.090 | 0.7590 | 1.000 |
| **Cholesterol** | 14 | -0.086 | 0.7710 | 1.000 |
| **Retinol** | 14 | -0.086 | 0.7710 | 1.000 |
| **Methionine** | 14 | 0.081 | 0.7820 | 1.000 |
| **Glutamic Acid** | 14 | 0.081 | 0.7820 | 1.000 |
| **SFA 17.0 margaric acid** | 14 | -0.077 | 0.7940 | 1.000 |
| **Lysine** | 14 | 0.077 | 0.7940 | 1.000 |
| **Arginine** | 14 | 0.077 | 0.7940 | 1.000 |
| **Tyrosine** | 14 | 0.072 | 0.8050 | 1.000 |
| **Sodium** | 14 | -0.072 | 0.8050 | 1.000 |
| **Total fat** | 14 | -0.068 | 0.8170 | 1.000 |
| **Threonine** | 14 | 0.064 | 0.8290 | 1.000 |
| **Isoleucine** | 14 | 0.064 | 0.8290 | 1.000 |
| **Alanine** | 14 | 0.064 | 0.8290 | 1.000 |
| **Serine** | 14 | 0.064 | 0.8290 | 1.000 |
| **MUFA 18.1 oleic acid** | 14 | -0.064 | 0.8290 | 1.000 |
| **Dietary Folate Equivalents** | 14 | -0.064 | 0.8290 | 1.000 |
| **PUFA 18.4 parinaric acid** | 14 | -0.052 | 0.8590 | 1.000 |
| **MUFA (total)** | 14 | -0.051 | 0.8640 | 1.000 |
| **CLA (trans-10, cis-12)** | 14 | 0.049 | 0.8690 | 1.000 |
| **PUFA (total)** | 14 | -0.046 | 0.8760 | 1.000 |
| **Total protein** | 14 | 0.046 | 0.8760 | 1.000 |
| **Leucine** | 14 | 0.042 | 0.8870 | 1.000 |
| **MUFA 22.1 erucic acid** | 14 | -0.040 | 0.8920 | 1.000 |
| **MUFA 14.1 myristoleic acid** | 14 | -0.037 | 0.8990 | 1.000 |
| **Stearic acid (SFA 18:0)** | 14 | 0.033 | 0.9110 | 1.000 |
| **Palmitic acid (SFA 16:0)** | 14 | -0.033 | 0.9110 | 1.000 |
| **Tagatose** | 14 | 0.032 | 0.9130 | 1.000 |
| **Cystine** | 14 | 0.029 | 0.9230 | 1.000 |
| **MUFA 20.1 gadoleic acid** | 14 | -0.029 | 0.9230 | 1.000 |
| **Aspartic Acid** | 14 | -0.029 | 0.9230 | 1.000 |
| **TRANS 18.2 trans octadecadienoic acid** | 14 | -0.029 | 0.9230 | 1.000 |
| **Phosphorus** | 14 | 0.024 | 0.9350 | 1.000 |
| **Choline** | 14 | -0.024 | 0.9350 | 1.000 |
| **Added Sugars by Available Carbohydrate** | 14 | -0.022 | 0.9400 | 1.000 |
| **TRANS 16.1 trans hexadecenoic acid** | 14 | -0.020 | 0.9460 | 1.000 |
| **SFA (total)** | 14 | -0.020 | 0.9460 | 1.000 |
| **PUFA 20.4 n 6 arachidonic acid .AA.** | 14 | 0.018 | 0.9520 | 1.000 |
| **Natural Folate food folate** | 14 | 0.011 | 0.9700 | 1.000 |
| **Total Folate** | 14 | -0.011 | 0.9700 | 1.000 |
| **Tryptophan** | 14 | 0.007 | 0.9820 | 1.000 |
| **SFA 22.0 behenic acid** | 14 | -0.004 | 0.9880 | 1.000 |
| **PUFA 18.2 linoleic acid. undifferentiated** | 14 | 0.002 | 0.9940 | 1.000 |
| **Animal protein** | 14 | -0.002 | 0.9940 | 1.000 |
| **PUFA 18.2 n 6 linoleic acid .LA.** | 14 | -0.002 | 0.9940 | 1.000 |
| **Omega 6 Fatty Acids** | 14 | -0.002 | 0.9940 | 1.000 |
| **PUFA 20.4 arachidonic acid. undifferentiated** | 14 | 0.000 | 1.000 | 1.000 |

**Supplementary Figure 1**

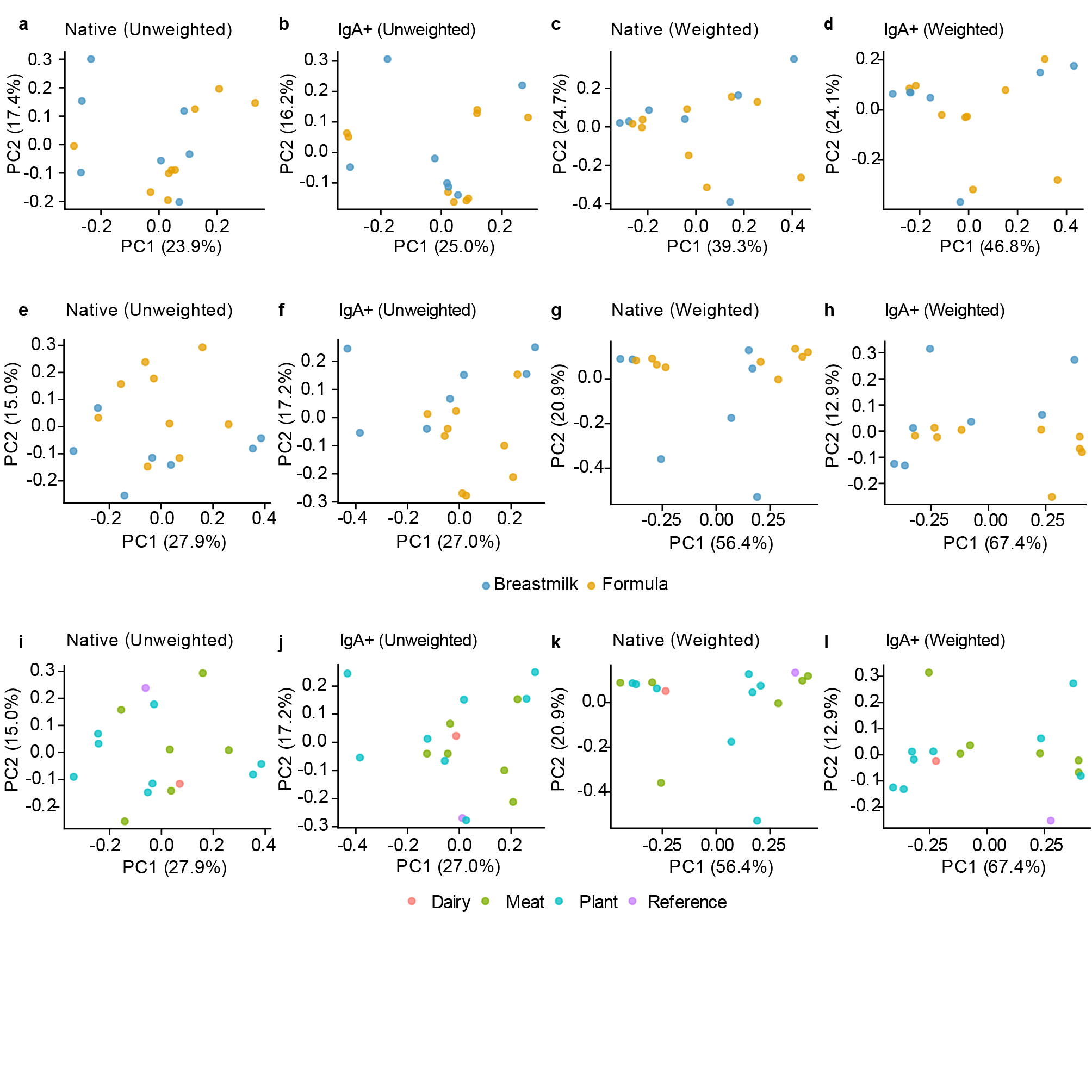

**Supplementary Figure 1. Community composition does not differ significantly by feeding mode or dietary intervention group. PCoA ordination of UniFrac distances for infant fecal samples.** **a-d**, 6-month (pre-intervention) samples colored by feeding mode (human milk vs Formula) for native unweighted (a), IgA+ unweighted (b), native weighted (c), and IgA+ weighted (d) UniFrac distances. n = 16 samples. e-h, 12-month (post-intervention) samples colored by feeding mode for the same four combinations. n = 16 samples. i-l, 12-month samples colored by dietary intervention group (Dairy, Meat, Plant, Reference). PERMANOVA revealed no significant separation by feeding mode at either timepoint or by dietary intervention group at 12 months (all P > 0.05; R² and P values reported in Supplementary Table S12). Percentage of variance explained by each principal coordinate axis is indicated on the axis labels.

**Supplementary Table S12.** PERMANOVA results for community composition by feeding mode and dietary intervention group (UniFrac distances, native and IgA-coated fractions).

| **Panel** | **Timepoint** | **Comparison** | **Distance** | **n** | **R2** | **F_stat** | **P** |
| --- | --- | --- | --- | --- | --- | --- | --- |
| **a** | 6mo | Feeding mode | Native (Unweighted) | 16 | 0.0686 | 1.03 | 0.424 |
| **b** | 6mo | Feeding mode | IgA+ (Unweighted) | 16 | 0.0685 | 1.03 | 0.368 |
| **c** | 6mo | Feeding mode | Native (Weighted) | 16 | 0.0282 | 0.41 | 0.863 |
| **d** | 6mo | Feeding mode | IgA+ (Weighted) | 16 | 0.0338 | 0.49 | 0.764 |
| **e** | 12mo | Feeding mode | Native (Unweighted) | 16 | 0.0714 | 1.08 | 0.344 |
| **f** | 12mo | Feeding mode | IgA+ (Unweighted) | 16 | 0.1 | 1.56 | 0.097 |
| **g** | 12mo | Feeding mode | Native (Weighted) | 16 | 0.0911 | 1.4 | 0.231 |
| **h** | 12mo | Feeding mode | IgA+ (Weighted) | 16 | 0.1119 | 1.76 | 0.167 |
| **i** | 12mo | Dietary intervention group | Native (Unweighted) | 16 | 0.1256 | 0.57 | 0.98 |
| **j** | 12mo | Dietary intervention group | IgA+ (Unweighted) | 16 | 0.1533 | 0.72 | 0.836 |
| **k** | 12mo | Dietary intervention group | Native (Weighted) | 16 | 0.1309 | 0.6 | 0.882 |
| **l** | 12mo | Dietary intervention group | IgA+ (Weighted) | 16 | 0.1808 | 0.88 | 0.578 |

**Supplementary Figure 2**

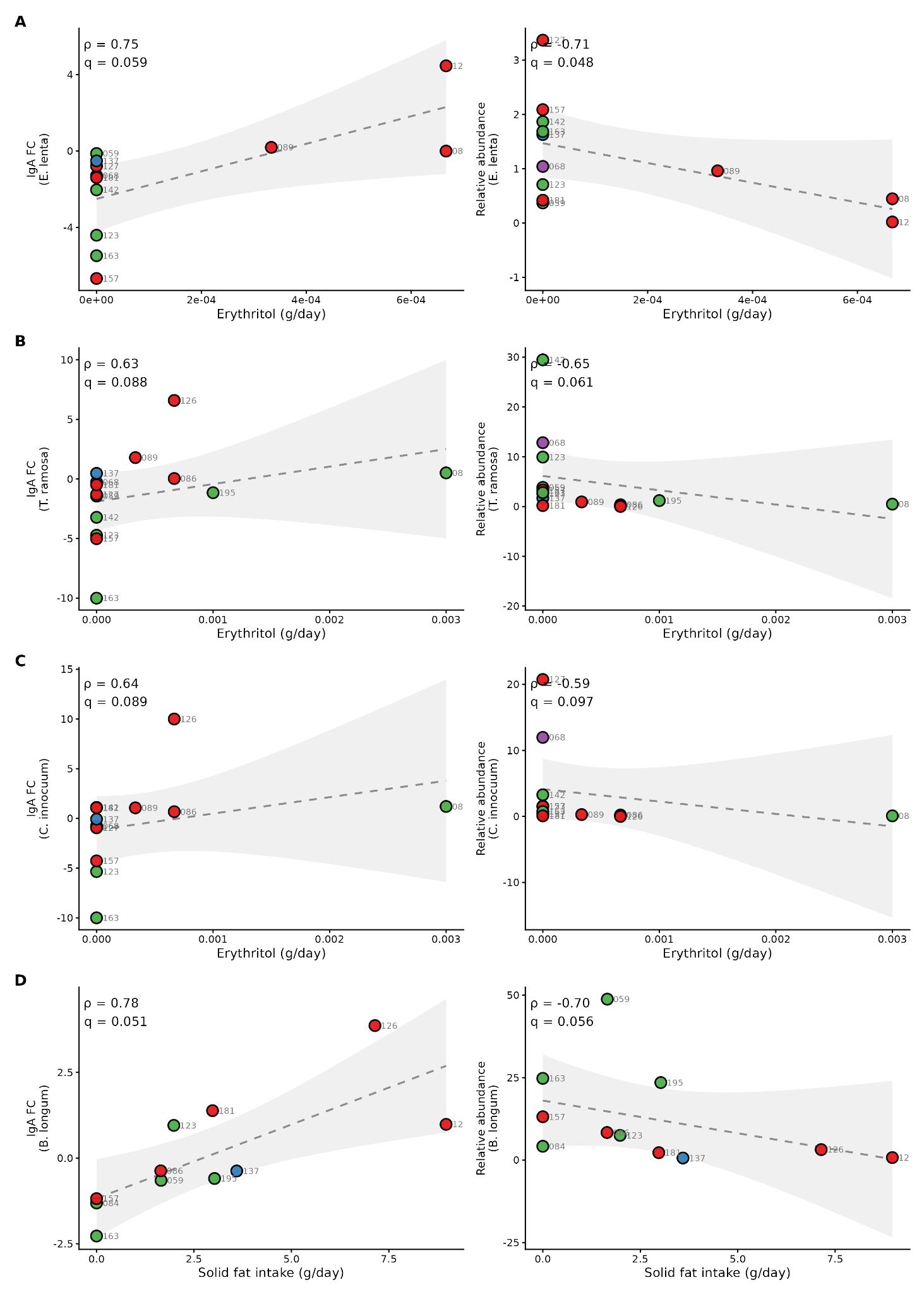

**Supplementary Figure S2.** Opposing associations between dietary nutrient intake and IgA targeting versus relative abundance for selected species. Spearman correlations between nutrient intake and species-level IgA fold change (FC; left column) or native relative abundance in the unsorted metagenomic fraction (right column) are shown for four species–nutrient pairs exhibiting divergent directionality: *Eggerthella lenta* versus erythritol **(A)**, *Thomasclavelia ramosa* versus erythritol **(B)**, *Clostridium AQ innocuum* versus erythritol **(C)**, and *Bifidobacterium longum* versus solid fat intake **(D)**. Each point represents one infant, colored by dietary arm. Dashed lines and shaded bands indicate the linear regression fit and 95% confidence interval. *ρ* and *q* values denote Spearman correlation coefficients and Benjamini–Hochberg-adjusted p-values, respectively.
